# Isolation of the first cultured representative from a deep-branching phylum in serpentinite-hosted ecosystems

**DOI:** 10.1101/2025.11.21.689855

**Authors:** Ge Liu, Jie Zhu, Ying Sun, Rui Liu, Yaru Gu, Shuai Yuan, Hongliang Wang, Mengna Li, Haibin Qi, Quancai Peng, Shimei Wu, Chaomin Sun

## Abstract

Serpentinization drives abiotic synthesis of organics (e.g., hydrocarbon) potentially conducive to the emergence of life, making serpentinite-hosted systems and associated microbial community key windows into nature of life’s origin. Although cultivation-independent studies uncovered the *Candisdatus* Bipolaricaulota widely distributed in serpentinizing environments, cultivation of this phylum has been unsuccessful. Here we cultured the first pure strain, J31, of *Ca.* Bipolaricaulota from the Lost City hydrothermal field, a well-characterized marine serpentinite system, using hydrocarbons as the primary nutrient source. As an early-branching bacterial lineage, strain J31 exhibits an unusual morphology composed of a central rod and elongated Toga-like extensions at both ends, and divides by binary fission. Strain J31 absorbs hexadecane through Toga ends via coordinated processes resembling inhalation and swallowing, after which hexadecane is efficiently transported to the central rod in the vesicle-like structures and subsequently converted into membrane lipids to support Toga synthesis and cellular growth. Hydrocarbon-degrading capability is widespread among the globally distributed members of *Ca.* Bipolaricaulota. Strain J31 colonizes serpentine minerals, facilitating the utilization of hydrocarbons derived from serpentines and promoting the release of soluble iron and silicon, thereby linking microbial activity to geochemical cycling. Thus, our study presents a novel strategy for cultivating deep-branching bacteria and offers insights into the metabolic foundations of early life on Earth—and potentially on other rocky planets undergoing serpentinization.

## Introduction

Microorganisms represent the most diverse and abundant cellular life forms on Earth, occupying every possible metabolic niche (*1*). Most remain uncultured (*2, 3*), and we have only recently become aware of their presence, metabolism and ecology mainly through omics-based cultivation-independent characterization (*4, 5*). Nevertheless, many microorganisms display physiological and biochemical traits that deviate substantially from metagenomic predictions, highlighting the urgent need for axenic cultures (*6, 7*).

Thermophiles are microorganisms that grow optimally above 45 °C (*8*), and they are found to be widely distributed in multiple artificial or natural high-temperature environments (*9*). Cultivation-independent analyses of hot springs in Yellowstone National Park reveal that geothermal systems host an abundance of uncultivated thermophiles with great phylogenetic and functional diversity, including deeply branching Archaea (*10*), and twelve novel lineages of Bacteria, provisionally named OP1-OP12 (*11*). Extreme conditions such as alkaline, acidic, high-temperature, and high-salinity environments are similar to that of the early Earth, and many geological studies and experimental evidences supported the early origin of thermophiles (*12, 13*). The “closely clustered thermophilic bacteria” (CCTB) first proposed in 2023 (*9*), refer to those deep-branching bacterial phyla that cluster closely near the root of the Archaea-Bacteria tree (*9*). Studying these early diverging thermophiles can shed light on the ecosystems of early Earth and adaptations of primitive life (*14, 15*). Yet, it is extremely difficult to obtain pure cultured novel early-branching thermophilic bacteria.

Deep-sea high-temperature environments where thermophiles inhabit are often associated with serpentinization, the hydration process of ultramafic rocks, accompanied by the production of serpentine minerals and hyperalkaline fluids rich in H2, CO, CH4 and various organic molecules (e.g., formate, methanol, and hydrocarbon) (*16–20*). These abiotic organic molecules may be potential prebiotic organic compounds for life’s emergence (*21*), and consequently, serpentinization-influenced environments are commonly regarded as one of the most plausible habitats for early life on Earth, as well as potential niches for life in other planetary systems where ultramafic rocks are present and capable of undergoing serpentinization, such as on Saturn’s icy moon Enceladus (*22*) or Mars (*23*). The serpentinite-hosted Lost City hydrothermal field (LCHF) is a remarkable submarine ecosystem where geological, chemical, and biological processes are intimately interconnected (*24*), with high-pH (9–11), 40–91 °C fluids rich in H2, CH4, and hydrocarbons (*24, 25*). Metagenomic and metatranscriptomic studies revealed the presence of diverse microbial communities, including sulfur-oxidizing/reducing, and methane-oxidizing bacteria, as well as methanogenic/methane-oxidizing archaea (*26–28*), in the serpentinite-hosted ecosystems. Metagenome-assembled genomes classified as *Methanosarcinaceae* and *Candidatus* Bipolaricaulota were recovered from venting fluids of LCHF, suggesting methanogenic and acetogenic metabolisms in this subseafloor ecosystem (*26*). Despite the potential for serpentinization to have fueled the metabolisms of life on early Earth, the specific adaptations that allow for life under these conditions remain poorly understood.

“*Ca.* Bipolaricaulota” (*29*) (formerly candidate division OP1 (*11*) and “*Ca.*Acetothermia”(*30*)) represents a deeply branching bacterial lineage (*30, 31*). Although metagenomic analyses documented its presence and potential lifestyle (*29*), the lack of cultured representatives hinders culture-based investigations into its metabolic capabilities (*32*) and ecological functions. Here, we report the isolation of the first cultured representative, strain J31, obtained from venting fluids of the serpentinite-hosted LCHF. We conduct a detailed investigation into its phylogeny, morphology, physiology, and ecology. This allows us, for the first time, to reveal an unusual biology of this unrecognized microbial player in the serpentinite-hosted hydrothermal vent systems.

## Results

### Cultivation of the first representative of *Ca.* Bipolaricaulota

Hydrocarbons serve as key electron donors for microorganisms inhabiting hydrothermal environments (*33*). Given the production of hydrocarbons during the serpentinization process, we developed a novel isolation and cultivation strategy, termed the “hydrocarbon utilization-driven” approach (Fig. 1B), to recover a greater diversity of microorganisms from venting fluids of the serpentinite-hosted LCHF (Fig. 1A). After being filtered and concentrated, venting fluid samples were immediately inoculated into an oligotrophic medium with crude oil, and incubated for one month at 65 °C for enrichment. A single-cell clone culture was then isolated using the dilution-to-extinction method. Following 3-5 consecutive transfers under optimal growth conditions and verification of purity via transmission electron microscopy (TEM) and 16S rRNA gene PCR, a pure culture, designated strain J31, was successfully obtained.

The 16S rRNA gene sequence of strain J31 confirmed its identity as the first cultured representative of the previously uncultivated *Ca.* Bipolaricaulota phylum. To resolve the definitive evolutionary position of this phylum, we constructed a robust phylogeny based on concatenated conserved proteins (Materials and Methods). This analysis conclusively places *Ca.* Bipolaricaulota within the CCTB supergroup (Fig. 1C), anchoring it within a key thermophilic assemblage that also includes phyla such as *Thermotogota*, *Thermodesulfobiota*, *Dictyoglomota*, *Caldisericota*, *Zhurongbacter*, and *Coprothermobacterota* (*9*). This finding is significant as it conclusively integrates a widespread, deep-branching, and previously uncultured phylum within the CCTB, substantially expanding the known phylogenetic breadth of this supergroup.

**Fig. 1.**
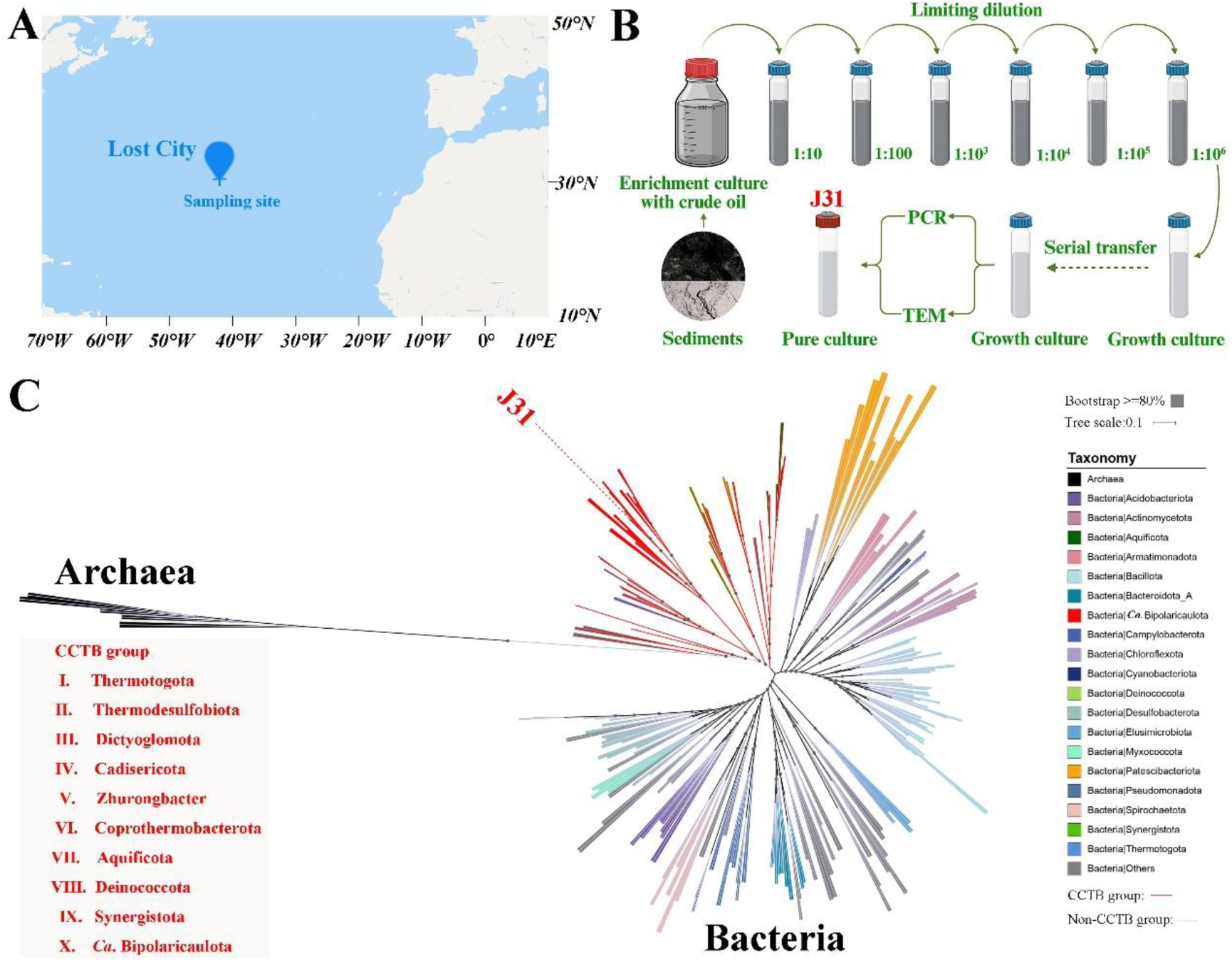
Isolation procedure and phylogenetic analysis of strain J31. (**A**) Location of the sampling site on the Lost City Hydrothermal Field (LCHF). (**B**) Schematic diagram of the isolation strategy used to isolate strain J31 in this study. (**C**) Phylogenomic tree of strain J31 and other major thermophilic bacteria (CCTB group) using Archaea as the outgroup with 33 conserved single-copy protein sequences (CSCP) with IQ-Tree using the -m MFP+MERGE model. The red branches highlight the CCTB group: *Thermotogota*, *Thermodesulfobiota*, *Dictyoglomota*, *Caldisericota*, *Zhurongbacter*, *Coprothermobacterota*, *Aquificota*, *Deinococcota*, *Synergistota* and *Ca*. Bipolaricaulota.

### Unique morphology and cell division mode of strain J31

Observations by phase-contrast microscopy (PCM) (Fig. 2A), laser confocal microscope (LSM) (Fig. 2B), scanning electron microscopy (SEM) (Fig. 2C), and TEM (Fig. 2D-F) show that the main bodies of strain J31 are rod-shaped (0.4 µm × 0.8-2 µm) with outer sheath-like structures (hereafter referred to as “Toga”, 0.2-0.3 µm × 0-10 µm) at both ends. Membrane staining with the fluorescent dye DiI demonstrates that these appendages are extensions of the cell envelope (Fig. 2B), similar to the Toga observed in members of the phylum *Thermotogota* (*34*) (Fig. S1). Based on the size of the central rod and the length or appearance of Toga, three different morphologies of strain J31 were observed (Fig. 2A-D): M1, central rod with two Toga structures of similar length; M2, smaller central rod with two Toga structures of different length; M3, smallest central rod with one single Toga structure. These different morphologies likely represent bacterial morphology at different growth stages. Cells of strain J31 are Gram-stained negative (Fig. S2), and the cytoplasm containing nucleic acids are mainly distributed in the rod-shaped “main body” (Fig. S3). Although typical Gram-negative cells only have two lipid bilayers (cytoplasmic membrane and outer membrane), cryo-EM revealed that the central rod of strain J31 exhibits an unusual multilayered cell envelope architecture with outer, inner, and intra-cytoplasmic lipid membrane-like layers (LMLs) (Fig. 2G), similar to another anaerobic, Gram-negative *Atribacterota* strain, *Atribacter laminatus* RT761 (*35*). Further observations using cryo-EM show that a proteinaceous surface layer (S-layer) is distributed on the outermost surface of the Toga, displaying the hexagonal pattern with 6.25 nm spacing (Fig. 2H and I).

**Fig. 2.**
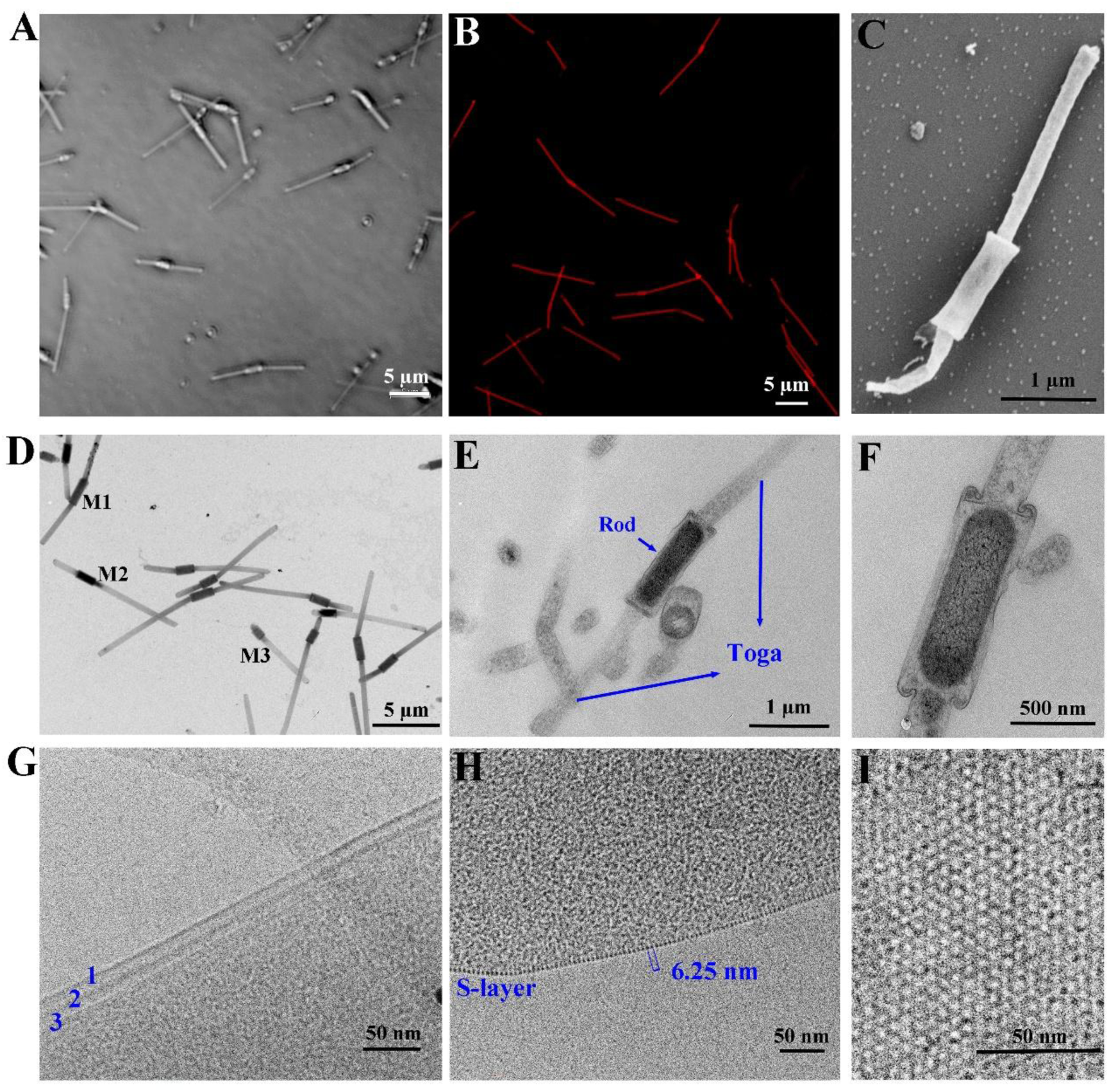
Morphology and membrane structure of strain J31. (**A**) Phase-contrast microscopy (PCM). (**B**) Laser confocal microscope (LSM) image after staining with the membrane dye DiI (red), demonstrating the Toga are extensions of the cell envelope. (**C**) Scanning electron microscopy (SEM). (**D** to **F**), Transmission electron microscopy (TEM) of a whole cell (D) and ultrathin sections (E, F). Labels M1-M3 indicate three distinct morphologies. (**G**) Cryo-EM image of the central rod showing the outer (1), inner (2), and intra-cytoplasmic (3) lipid membrane-like layers (LMLs). (**H** and **I**) Cryo-EM side (H) and top (I) views of the Toga S-layer, revealing a hexagonal pattern with 6.25 nm spacing.

To investigate the mode of cell division, strain J31 in the exponential stage was observed by TEM. As shown in Fig. S4, different morphologies of strain J31 appear within the same field of view, and shapes from I to X collectively represent almost the entire cell division cycle: small central rods with a single Toga represent cells just after cell division (shape I), and the longer rods with two Toga of equal length represent cells just before cell division (shape X). Further observations demonstrate that strain J31 reproduces via typical binary fission (Fig. 3A-C). Prior to division, both the central rod and the Toga elongate along their longitudinal axis, nearly doubling in size (Fig. 3A, I-V). Key cellular components in the central rod, including DNA and ribosomes, segregate to form two distinct compartments, followed by the formation of a septum at the middle of cells (Fig. 3C, I-VI). The septum then continues to extend until it splits entirely, and the two daughter cells separate (Fig. 3A, VI-VIII and Fig. 3B, I-V).

**Fig. 3.**
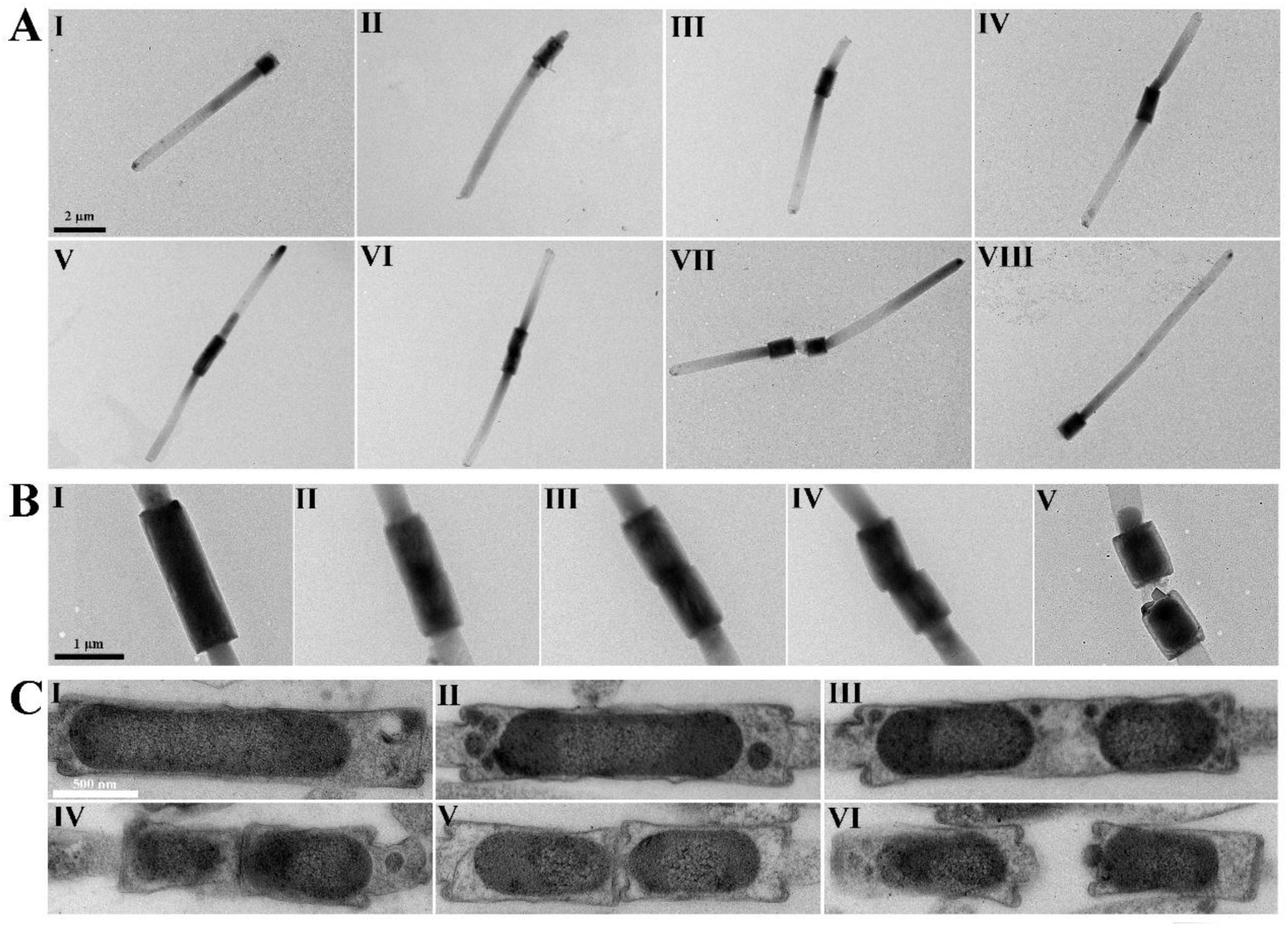
Binary fission mode of strain J31. (**A**) TEM images of strain J31 at different growth stages. Panels I–VIII represent nearly the entire cell division cycle of strain J31. (**B**) TEM images illustrating the morphological changes of the central rod during division (Panels I–V). (**C**) TEM images of ultrathin sections showing internal septum formation and division process (Panels I–VI).

### Efficient uptake, transport, and metabolism of hydrocarbons by strain J31

Since strain J31 was enriched and isolated from a hydrocarbon-containing medium, we next investigated its hydrocarbon metabolism. To this end, strain J31 was cultured in OM medium supplemented with hexadecane for 10 days. Cells grown with hexadecane (67 mg/L) presented a faster growth rate and higher biomass than those grown without hexadecane (Fig. 4A). Meanwhile, concomitant degradation of hexadecane by strain J31 was quantified using gas chromatography (Fig. 4A). It was evident that hexadecane removal rates increased progressively over time, with the maximum degradation rate reaching 47%. To visualize the utilization of hexadecane, FITC-labeled hexadecane (C16-FITC) was synthesized and added to the culture of strain J31. As shown in Fig. 4B and Movie S1, C16-FITC was rapidly absorbed by strain J31, and most of the green fluorescence signals emitted by C16-FITC were distributed in the central rods, indicating that the utilization and metabolism of hexadecane likely occur in the cytoplasm of strain J31. Consistently, observations by colloid gold immunoelectron microscopy demonstrated that hexadecane was primarily located in the main body of strain J31 (Fig. 4C). Following a 4-h incubation with hexadecane, transcriptomic analysis revealed a clear upregulation of genes involved in fatty acid and membrane lipid biosynthesis (Fig. 4D and E), suggesting that hexadecane supports the rapid growth of strain J31 by being converted into cellular lipids, important structural components of the Toga. To further investigate the metabolism and bioconversion of hexadecane in strain J31, ^13^C-labeled hexadecane (hexadecane-1,2-^13^C2) was added to the OM medium, and its intracellular flow was analyzed using the stable isotope technique via non-target metabolomics. The results showed that hexadecane was mainly metabolized into free fatty acids and membrane lipids to support cell growth, with membrane lipids exhibiting a higher ^13^C-labeling level than fatty acids (Table S1).

**Fig. 4.**
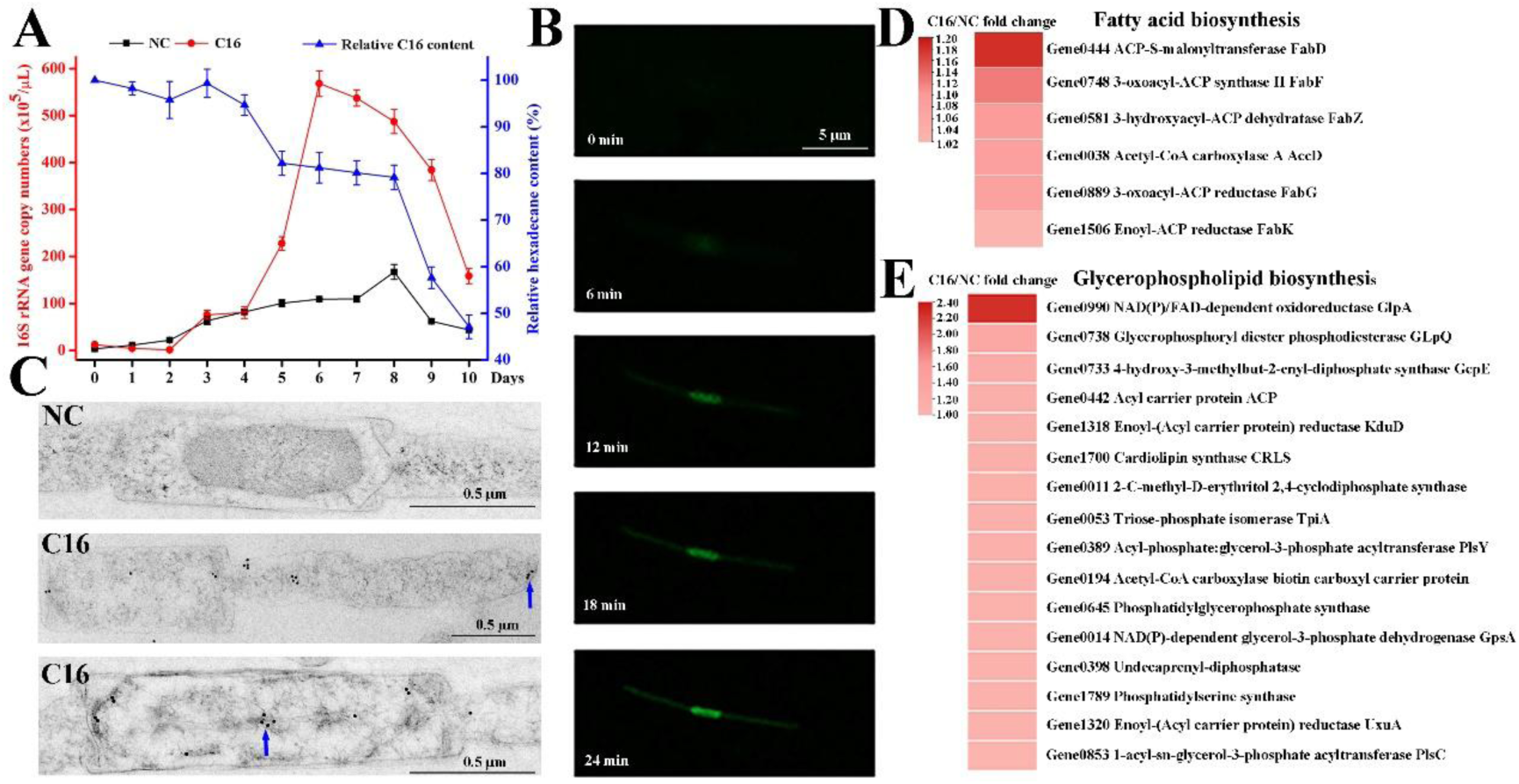
Uptake, transport, and metabolism of hexadecane (C16) by strain J31. (**A**) Strain J31 degraded C16 to promote its growth. Black and red lines represent the growth curves without and with C16, respectively. The blue line represents the relative content of C16. Cell growth was determined based on 16S rRNA gene copy number. (**B**) Fluorescence micrographs showing rapid uptake of FITC-labeled hexadecane (C16-FITC, green fluorescence) by strain J31. (**C**) Colloidal gold immunoelectron microscopy revealing C16 was mainly localized in the central rods. The small black dots indicated by the arrows are the signals of colloid gold particles. (**D** and **E**) Transcriptome analysis showing that C16 significantly upregulated the biosynthesis of both fatty acids (D) and membrane lipids (E).

**Movie S1.** The video shows the rapid uptake of FITC-labeled hexadecane (C16-FITC) by strain J31. Cells were incubated with C16-FITC and immediately imaged using a confocal laser scanning microscope. Images were acquired at one-minute intervals over a period of 1 h.

As displayed in Movie S1, the green fluorescence signals emitted by C16-FITC were translocating from the Toga ends to the central rod of strain J31 in a granular pattern. To explore the transport mode of hexadecane in detail, strain J31 incubated with C16-FITC was further observed by LSM. The granular green fluorescence signals migrated directionally from the Toga ends to the central rod (Fig. 5A). Co-localization of C16-FITC with the membrane dye DiI indicated that hexadecane was transported via vesicle-like structures in strain J31 (Fig. 5B). Micrographs of ultrathin sections confirmed the presence of vesicle-like structures in strain J31 (Fig. 5C). These vesicle-like structures are ultimately transported into the central rods, where they undergo membrane fusion with the cytoplasmic membrane and released their hexadecane cargo into the cytoplasm (Fig. 5C). When incubated with crude oil for 1 h, dark vesicle-like structures were again observed, forming at and detaching from the Toga ends (Fig. 5D and Fig. S5). Consistent with this, observations of ultrathin sections revealed strain J31 absorbed crude oil at the Toga ends, and finally transported it to the central rods in the vesicle-like structures (Fig. 5E). Cryo-EM imaging provided detailed insights into the architecture of the Toga ends (Fig. 5F, Figs. S6-7). Distinct structural states were observed, composed of three major components: the cytostome-like (1), cytopharynx-like (2), and zipper-like (3) structures, with a few pili distributed around the cytostome-like region. Based on these observations, we propose a model for absorption of hexadecane by strain J31 (Fig. 5G). The zipper-like structure first opens, allowing the cytostome-like structure to invaginate and draw hexadecane into the cytopharynx-like compartment with the help of pili; subsequently, the zipper-like structure closes, restoring the plump morphology of the Toga ends, while the hexadecane is encapsulated into vesicle-like structures and transported toward the central rod of strain J31. The cyclic repetition of these steps ensures efficient hexadecane uptake and intracellular transport.

**Fig. 5.**
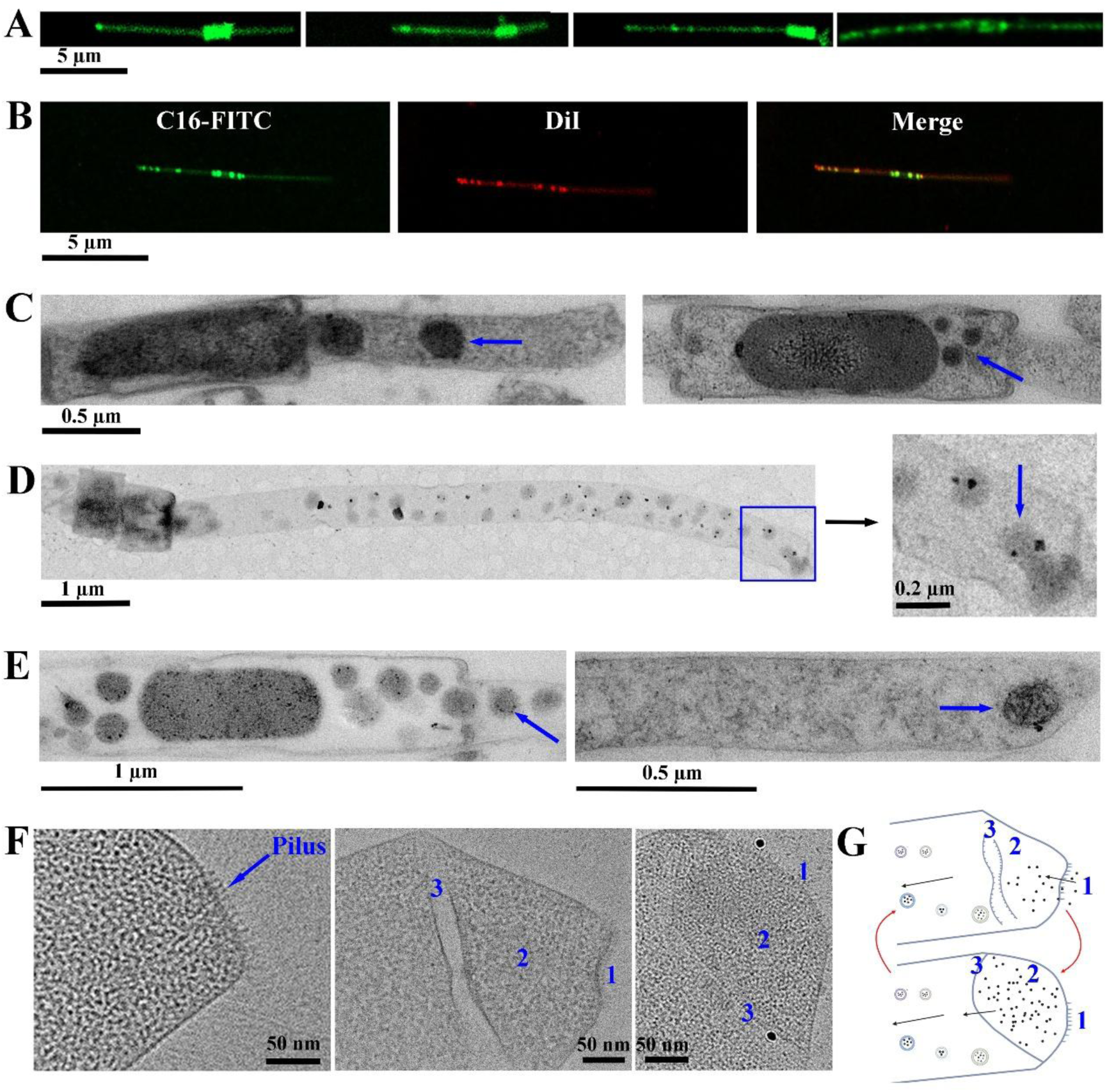
Vesicle-like transport of hexadecane (C16) by strain J31. (**A**) Vesicle-like transport of FITC-labeled hexadecane (C16-FITC) observed by LSM. (**B**) Co-localization of C16-FITC and the membrane dye DiI. (**C**) TEM images of ultrathin sections showing vesicle-like structures (blue arrow). The panels represent in the Toga (left) and the central rod (right). (**D**) TEM images showing vesicle-like structures of crude oil forming at the Toga ends. Enlarged view of the boxed regions is shown on the right. The blue arrow indicates the vesicle formation process. (**E**) TEM images of ultrathin sections showing crude oil-filled vesicle-like structures (blue arrow) in the central rod (left) and forming at the Toga end (right). (**F**) Cryo-EM images revealing the ultrastructure of the Toga ends. The left panel shows pili at the ends of Toga. The middle and right panels show the cytostome-like (1), cytopharynx-like (2) and zipper-like (3) structures. (**G**) Proposed model for C16 transport. Hexadecane is absorbed through coordinated actions of three organelle-like structures: the cytostome-like (1), cytopharynx-like (2) and zipper-like (3). Black arrows indicate the transport pathway, and red arrows indicate the conversion of different states.

### Global distribution and hydrocarbon metabolism of *Ca.* Bipolaricaulota

To elucidate the ecological niches of *Ca*. Bipolaricaulota, ninety-nine 16S rRNA gene sequences were retrieved from the NCBI database (Table S2), and their geographic locations and isolation sources were analyzed (Fig. S8A). Collectively, *Ca.* Bipolaricaulota appears to have a global distribution, and anaerobic environments are common to most of these habitats, e.g., hot springs, hydrothermal vents, oil reservoirs, saline lakes, wastewater treatment plants, groundwater, continental subsurface, tailing ponds, marine animal tissues, deep-sea sediments, cold seeps, landfills, Antarctica and human stool. This broad distribution indicates that members of this phylum possess diverse physiological capabilities and play important ecological roles.

To further study the hydrocarbon degradation capability of *Ca.* Bipolaricaulota, 189 complete and incomplete genomes (MAGs, metagenome-assembled genomes) were obtained from the NCBI database (Table S3). These genomes were aligned against the CANT-HYD database to identify genes involved in anaerobic and aerobic degradation pathways of aliphatic and aromatic hydrocarbons. 76 out of the 189 genomes encoded key hydrocarbon degradation genes, accounting for approximately 40.2% (Table S4). As shown in Fig. S8B, enzymes involved in anaerobic hydrocarbon degradation (AssA, BssA, NmsA, AhyA, CmdA, and EbdA) exhibited a relatively widespread distribution across these genomes, indicating a strong potential for anaerobic degradation of aliphatic and aromatic hydrocarbons. DszC, CYP153, LadA-alpha, LadA-beta, l LadB, non-NdoB, AbcA-1/2, and K27540, responsible for the aerobic hydrocarbon degradation, were also annotated in few genomes, consistent with the predominantly anaerobic lifestyle of this phylum. Collectively, these findings highlight the important ecological role for members of *Ca.* Bipolaricaulota in the global carbon cycle.

### Contribution of strain J31 to the elemental cycle in serpentinite-hosted ecosystems

Abiotic hydrocarbons production is considered to be driven by serpentinization, and laboratory experiments have demonstrated that hydrocarbons with up to 24 carbon atoms can be synthesized through hydrothermal reduction of NaHCO3 (*36*). To determine whether serpentinites contain hydrocarbons, *n*-alkanes were extracted from serpentine minerals, which were confirmed to be serpentinite through cross-polarized microscopy and elemental composition analyses (Fig. S9, Fig. S10A). As shown in Table S5, straight-chain alkanes (*n*-alkanes) with carbon chain lengths ranging from C11 to C30 were identified, suggesting that these *n*-alkanes are encapsulated within serpentine minerals. Among them, hexadecane exhibits the highest content, reaching 11.70 ± 1.24 ng/g of serpentine minerals. As displayed in Fig. 4A, hexadecane markedly promoted the rapid growth of strain J31, indicating that microbially mediated utilization of abiotic hydrocarbons may play a role in linking geochemical and biological processes in serpentinizing environments.

To further study the specific metabolic interactions of strain J31 within the serpentinite-hosted systems, cells were incubated with deep-sea serpentine minerals in the OM medium for 3 weeks. As observed by SEM, strain J31 exhibited tight colonization on serpentine minerals, with some individuals having infiltrated into the rocks’ fractures (Fig. 6A), indicating that serpentine-derived conditions are compatible with its growth. In the process of co-cultivation with serpentine minerals, strain J31 promoted the formation of black precipitates (Fig. 6B). Further characterization using energy-dispersive spectroscopy (EDS) confirmed these precipitates to be composed of sulfur, iron, silicon, and magnesium (Fig. S10B), consistent with the elemental composition of serpentine minerals used (Fig. S10A). Supernatants from the co-cultures were collected and then analyzed using an ICP-OES to monitor changes in the concentrations of soluble iron, silicon, and magnesium. As the co-cultivation time prolonged, serpentine minerals spontaneously released iron and silicon into the culture system, whereas strain J31 enhanced this release (Fig. 6C and D). Iron is a vital cofactor in numerous metabolic processes across all life forms (*37*), while dissolved silicon serves as an essential nutrient for deep-sea hexactinellid sponges (*38*). Thus, the bio-mediated release of soluble iron and silicon by strain J31 may provide critical resources for other vent-associated organisms, contributing to iron and silicon cycling in the deep-sea ecosystem. Magnesium release, however, was not detected—likely due to the already high Mg^2+^ concentration in seawater (Fig. S11).

**Fig. 6.**
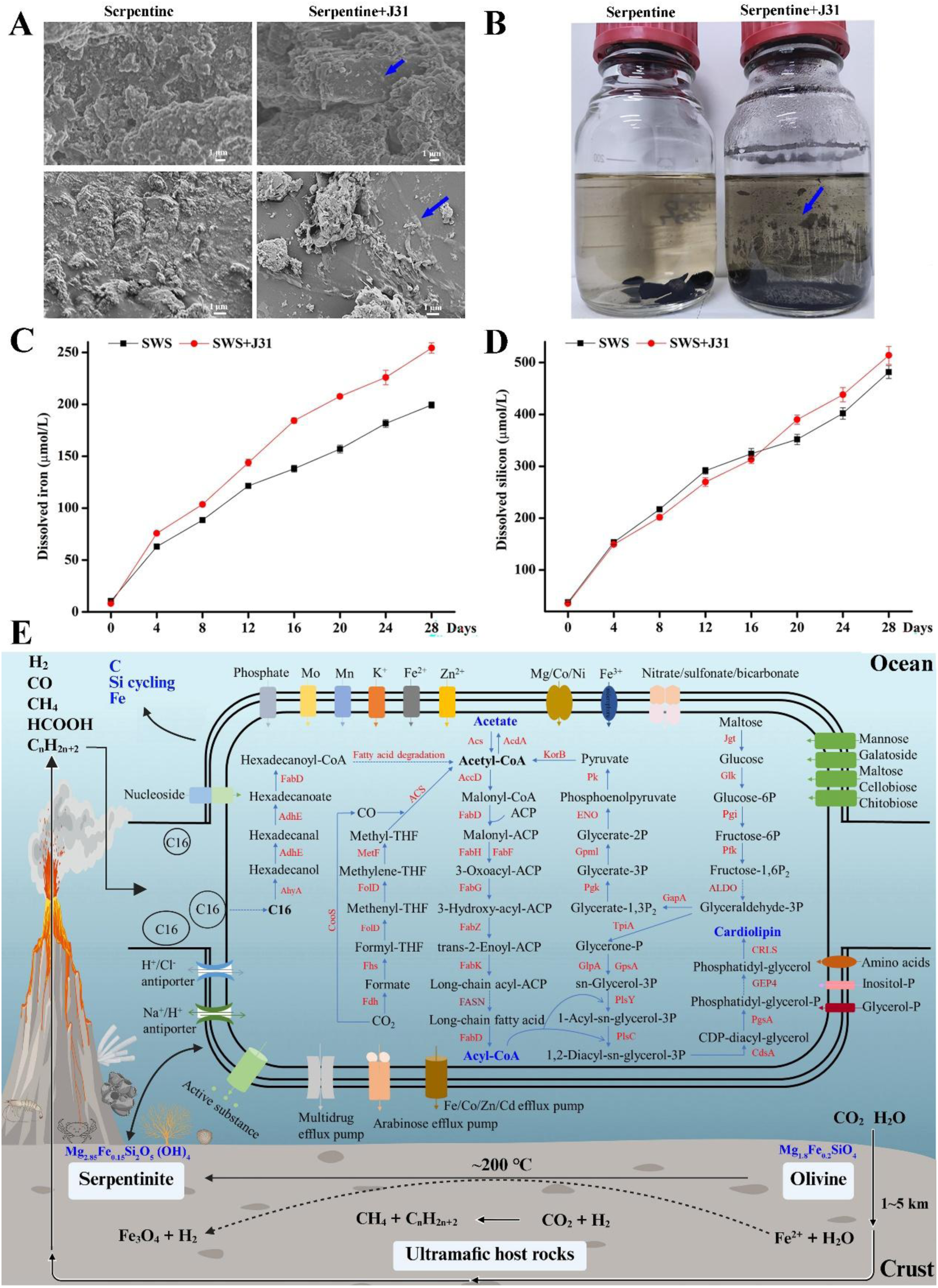
The interaction between strain J31 and serpentinite minerals drives the elemental cycle in deep-sea serpentinite-hosted ecosystems. (**A**) SEM images showing colonization of strain J31 on serpentine minerals. The blue arrows indicate the positions of strain J31. (**B**) Co-cultivation of strain J31 with deep-sea serpentine minerals induced the formation of black precipitates (indicated by the blue arrows). (**C** and **D**) Strain J31 promoted the dissolution and release of iron (C) and silicon (D) from serpentine minerals, as measured by inductively coupled plasma optical emission spectroscopy (ICP-OES). SWS refers to serpentine minerals. (**E**) Proposed metabolic model of strain J31 in deep-sea serpentinite-hosted environments. Serpentinization in the LCHF produces various abiotic organic molecules. Inhabiting this ecosystem, strain J31 absorbs and utilizes hexadecane to support its growth, accompanied by dissolution of serpentinite minerals and release of Si and Fe. Abbreviations: C16, hexadecane; AhyA, molybdopterin-family alkane C2 methylene hydroxylase; AdhE, bifunctional aldehyde-alcohol dehydrogenase; FabD, ACP-S-malonyltransferase; ACS, CO dehydrogenase/acetyl-CoA synthase; MetF, methylenetetrahydrofolate reductase; FolD, 5,10-methylene-tetrahydrofolate dehydrogenase; Fhs, formate-tetrahydrofolate ligase; Fdh, formate dehydrogenase; AcdA, acetyl-CoA decarbonylase/synthase; AccD, acetyl-CoA carboxylase; FabH, 3-oxoacyl-ACP synthase; FabF, 3-oxoacyl-ACP synthase II; FabG, 3-oxoacyl-ACP reductase; FabZ, 3-hydroxyacyl-ACP dehydratase; FabK, enoyl-ACP reductase; KorB, 2-oxoglutarate/2-oxoacid ferredoxin oxidoreductase; Pk, pyruvate kinase; ENO, enolase; GpmI, phosphoglycerate mutase; Pgk, phosphoglycerate kinase; GapA, glyceraldehyde-3-phosphate dehydrogenase; Pfk, ATP-dependent 6-phosphofructokinase; Pgi, glucose/mannose-6-phosphate isomerase; Glk, glucokinase; Jgt, alpha-amylase/alpha-mannosidase; TpiA, triose-phosphate isomerase; GlpA, NAD(P)/FAD-dependent oxidoreductase; GpsA, NAD(P)-dependent glycerol-3-phosphate dehydrogenase; PlsY, acyl-phosphate:glycerol-3-phosphate acyltransferase; PlsC, 1-acyl-sn-glycerol-3-phosphate acyltransferase; CdsA, phosphatidate cytidylyltransferase; PgsA, CDP-diacylglycerol-glycerol-3-phosphate 3-phosphatidyltransferase; CRLS, cardiolipin synthase.

To illustrate the ecological roles of strain J31 in the serpentine-hosted environments, its metabolic model was constructed based on genomic annotations, experimental evidences and geological studies (Fig. 6E). Serpentinization acts as a planetary scale reducing agent (*39*). During serpentinization in the LCHF, seawater infiltrates the crust cracks where it reacts with ultramafic rocks at temperatures of approximately 200 °C. In this reaction, Fe^2+^ in olivine is oxidized by water to Fe^3+^, generating iron oxides in serpentinite, as well as magnetite (Fe3O4), and H2. The resulting reducing environment further promotes the abiotic conversion of CO2 to formate, methane, hydrocarbon and other organics. Within this setting, strain J31 absorbs hydrocarbons (taking hexadecane as an example) through its Toga ends and transports hexadecane via vesicle-like structures to the central rod, where hexadecane is converted into C16-CoA and ultimately degraded into acetyl-CoA. The acetyl-CoA is subsequently used for the synthesis of fatty acids and membrane lipids, supporting rapid cell growth. Inhabiting active serpentinization zones, strain J31 not only utilizes serpentinite-derived hydrocarbons but also facilitates the release of iron and silicon from serpentinite minerals—nutrients that can be assimilated by other members of the hydrothermal community. Collectively, these findings suggest that members of *Ca.* Bipolaricaulota play integral roles in the coupled cycling of carbon, iron, and silicon in serpentinite-hosted ecosystems, linking microbial metabolism to geochemical transformations in the deep sea.

## Discussion

In this study, we propose a “hydrocarbon utilization-driven” strategy to achieve the isolation of uncultured or difficult-to-culture species inhabiting venting fluids of the serpentinite-hosted LCHF. Hydrothermal activity generates numerous and diverse hydrocarbon compounds, thereby fostering the unique hydrocarbon-utilizing microbial communities (*33*). Hydrocarbons and high temperature are the two most critical environmental factors for these venting microorganisms. The successful cultivation of the first pure culture belonging to the *Ca*. Bipolaricaulota has shown that the culturing of uncultured or rare bacteria is indeed possible when their ecological niches are sufficiently well mimicked. The application of similar cultivation strategies to other understudied microorganisms will increase both the number and phylogenetic diversity of axenic cultures in the future.

The origin of organic matter remains a fundamental problem in the natural sciences even in the 21^st^ century. The discovery of hydrothermal activity in the late 1970s provided a new impetus for exploring the genesis and nature of organic compounds in the geological environment. Unsurprisingly, many theories on the origin of life invoke the geological process of serpentinization in hydrothermal vents, as its products are potential precursors for the emergence of early life (*40*). In addition to biogenic hydrocarbons, recent studies suggest that the abiogenic contribution to sources of hydrocarbons in hydrothermal fields should not be ignored (*19, 20*). This deep and hot biosphere may be partially energetically supported by hydrocarbons (*41*); however, we know little about this unique ecosystem. Distinct extremophiles inhabiting deep-sea hydrothermal vents may mediate the acquisition of deep-subsurface matter and energy, thereby providing clues for understanding the coupling between deep-sea—or even potentially early—life and the abiotic processes occurring beneath the seafloor. As an early-branching extremophile inhabiting the serpentinite-hosted LCHF, strain J31 colonizes serpentinite minerals, utilizes trapped hexadecane derived from serpentinites to fuel its growth, and simultaneously enhances the release of iron and silicon from the serpentinite minerals, thereby providing nutrients for other venting organisms such as deep-sea sponges. These activities suggest that the cycling of carbon, iron, and silicon in this marine serpentinite system is driven, at least in part, by strain J31.

The cell membrane is one of the most ancient and essential cellular components and is traditionally divided into Gram-positive (monoderm) and Gram-negative (diderm) (*42*). Recent phylogenomic analyses support the hypothesis that the last universal common ancestor (LUCA) possessed a diderm envelope (*43*), and that the outer membrane (OM) was subsequently lost multiple times during evolution, giving rise to monoderms. *Thermotoga* represents an early-branching bacterial phylum often placed near the root of the tree of life and therefore provides valuable insights into the nature of LUCA. The most well-known of *Thermotoga* is that all isolated members possess an OM detached at the cell poles (*44*), a structure commonly referred to as “Toga”. A similar Toga was recently observed both in the first enriched member (strain Ran-1) (*29*) and the first pure culture (strain J31) of the *Ca*. Bipolaricaulota. Interestingly, these two bacterial phyla are phylogenetically closely related. These unique OM detachments have been proposed to accommodate space for the degradation of organic matter in the periplasm in the case of *Thermotoga* (*45*), and to serve as specialized channels for hydrocarbon transport in the case of strain J31. Some diderms display intra-cytoplasmic membranes. A well-known example is *Atribacterium laminatus* RT761 (*35*), the first isolate of *Atribacterota*, which revealed an unusual diderm architecture under cryo-electron tomography, consisting of a cytoplasmic membrane (CM), an OM, and an intra-cytoplasmic membrane surrounding the nucleoid (*35*). Similarly, the central rod of strain J31 exhibits a three-layered cell envelope architecture with outer, inner, and intra-cytoplasmic lipid membranes. It is hypothesized that the primordial cell had only one membrane, it remains unknown how the second membrane could have evolved (*43*). The mechanisms underlying the biogenesis of these additional intra-cytoplasmic membranes may fundamentally reshape our view of cell envelope transitions. Further isolation and characterization of novel bacterial phyla are likely to yield important insights into the diversity and evolutionary transitions of cell envelope structures among previously uncultured microorganisms.

By combining TEM with cryo-EM, we observed unique ultrastructural features of the Toga ends of strain J31, including cytostome-like, cytopharynx-like, and zipper-like structures. These organelle-like features are likely to modulate the internal gas pressure within the Toga through coordinated actions such as inhalation, swallowing, and rhythmic opening/closing, thereby facilitating the rapid uptake of hexadecane. The absorbed hexadecane is subsequently transported from the Toga ends to the central rod via vesicle-like structures and ultimately utilized within the cytoplasm. This study provides rare evidence that bacteria may acquire organic nutrients through a pinocytosis-like mechanism, reminiscent of the feeding behavior observed in ciliates. This discovery reveals a distinctive feature of the *Ca*. Bipolaricaulota—simple in organization yet remarkably sophisticated, primitive yet functionally advanced. More efforts are still needed in the future to further understand the specific details and possible regulation mechanisms.

Most lineages within the domains Bacteria and Archaea remain uncharacterized due to limitations in sampling and cultivation. These difficult-to-culture (or uncultured) microorganisms, often referred to as the “dark matter” of life (*1*), are increasingly recognized as untapped ‘talented’ producers of small molecules with a potentially therapeutic character. Through genome analysis of strain J31, we identified a complete gene cluster responsible for terpenoid biosynthesis, and successfully isolated one active terpenoid alkaloid-like compound exhibiting potent antibacterial activity (Fig. S12). Generally, bacteria with large genomes, are considered ideal candidates for the production of novel antibiotics by hitherto unknown biosynthetic pathways (*6*). Remarkably, the fact that strain J31—with its relatively small genome—can still encode bioactive metabolites substantially expands our understanding of the metabolic potential of microorganisms.

Members of *Ca.* Bipolaricaulota are globally distributed in various habitats, particularly the hot and anaerobic environments. Strain J31 is a thermophilic, anaerobic and heterotrophic bacterium, isolated from the venting fluids of the serpentinite-hosted LCHF. Phylogenetic analysis indicates the early-branching of strain J31 that is often placed near the root of the tree of life. Interestingly, strain J31 exhibits an unusual morphology consisting of a central rod flanked by Toga-like structures at both ends. It is the first time this type of morphology has been reported, which can shed new light on the evolution of cell morphology. Moreover, strain J31 transports and utilizes hexadecane, a potential product of serpentinization, in the vesicle-like structures, thereby enabling rapid metabolism and bioconversion. In the serpentine-hosted environment, strain J31 also accelerates the dissolution of serpentine minerals and the release of soluble iron and silicon, consequently enhancing the biogeochemical cycling of these elements in the deep sea. Taken together, our findings contribute to a broader interdisciplinary understanding of how serpentinization-associated processes can support and shape early life on Earth or even on other rocky planets undergoing serpentinization.

### Taxonomic proposal to rename *Ca.* Bipolaricaulota to *Thermolongumtogota*

The name *Bipolaricaulota* refers to the presence of stalks at both poles of the rod-shaped cells. However, cells with two stalks at both poles correspond to the pre-division stage, whereas those with only a single stalk represent mature cells. *Ca*. Bipolaricaulota is closely related to *Thermotogota* (Fig. 1), both positioned near the root of the tree of life. The stalks of *Ca*. Bipolaricaulota are extensions of the cell envelope (Fig. 2B), analogous to the Toga observed in *Thermotogota* (Fig. S1). Given their close evolutionary relationship and morphological similarity between *Ca*. Bipolaricaulota and *Thermotogota*, we propose renaming the *Ca*. Bipolaricaulota as *Thermolongumtogota*, with stain J31 representing its first cultured member.

### Description of *Thermolongumtoga* gen. nov

*Thermolongumtoga* (Ther.mo.lon.gum.to’ga. Gr. fem. n. thermê, heat; L. masc. adj. longus, long; L. fem. n. toga, a toga (a loose outer garment); N.L. fem. n. *Thermolongumtoga*, a long, thermophilic bacterium with a Toga-like structure, referring to its thermophilic property, long cell shape, and the sheath-like outer membrane).

Obligately anaerobic, Gram-negative, non-motile, non-spore forming, rod cells accompanied with an outer sheath-like structure at both ends of the cell. Anaerobic respirations with alkane and sulfate are observed. The cellular fatty acids are C16:0 (44.63% of total), C14:0 (24.14%), C18:0 (7.49%), anteiso-C15:0 (6.07%), anteiso-C17:0 (3.93%), C12:0 (2.47%), C10:0 (2.08%), and C18:1 w9c (2.69%). DNA G+C content of the type species is 64.5 mol%. The type species is *Thermolongumtoga alcanivorasis*.

### Description of Thermolongumtoga alcanivorasis sp. nov

*Thermolongumtoga alcanivorasis* (al.ca.ni.vo’ra.sis. N.L. neut. n. alcanum, alkane (hydrocarbon); L. part. adj. vorans, devouring, eating; N.L. masc. adj. alcanivorasis, alkane-devouring, referring to the bacterium’s ability to utilize alkanes as a carbon and energy source).

Shows the following characteristics in addition to those given for the genus. Cells are Gram-negative rods (0.4 µm wide and 0.8∼2.0 µm long) with a small sheath-like structure at both ends (0.2∼0.3 μm wide and 0∼10.0 μm long). Grows at temperature range of 45–65 °C (optimal 55 °C), NaCl range of 1.0–3.0% (w/v) (optimal 2.0%), pH range of 6.0–8.0 (optimal pH 7.0). Yeast extract, tryptone and hexadecane supported the rapid growth. Growth occurs with maltose, lactose, trehalose, galactose, gelatin, pectin, and sodium sulfate. Cellulose, laminarin, fucoidan, xylan, sodium nitrate, sodium acetate, glucose, rhamnose, xylose, and arabinose are not utilized. Sensitive to kanamycin, gentamicin, chloramphenicol, and hygromycin, but resistant to ampicillin, rifampin, lovastatin, and simvastatin. No colonies were formed on agarose and gellan slants under the tested conditions. The type strain, J31^T^ (= MCCC 1K10091^T^), was isolated from venting fluids of Lost City Hydrothermal Field.

The complete genome of strain J31 is a single circular chromosome of 1,887,171 bp with a GC content of 64.5 mol% (Fig. S13). It contains three rRNA gene operons and 45 tRNA genes. A total of 1,837 coding DNA sequences (CDSs) with an average length of 949.16 bp, covering 92.39% of the whole genome, can be identified. Consistent with previous genomic analyses of other genomes and MAGs from “*Ca.* Bipolaricaulota” (*29, 46*), genes encoding АВС-type transporters responsible for uptake of sugars, amino acids, glycerol-3-phosphate, and peptides into the cell are detected, indicating that strain J31 can take up these compounds for use as carbon and energy sources. The genome encodes an incomplete tricarboxylic acid pathway, with only fumarate hydratase and succinyl-CoA synthetase annotated. Genome analysis reveals all enzymes of glycolysis are present, suggesting that imported sugars are likely to be metabolized via the Embden-Meyerhof pathway. The complete Wood-Ljungdahl (WL) pathway is identified in the strain J31 genome and other *Ca*. Bipolaricaulota genomes(*32*), indicating its ubiquitous phylum-wide distribution.

### Description of *Thermolongumtogaceae* fam. nov

*Thermolongumtogaceae*(Ther.mo.lon.gum.to.ga.ce’ae.N.L. fem. n. *Thermolongumtoga*, type genus of the family; suff. -aceae, ending to denote a family; N.L. fem. pl. n. *Thermolongumtogaceae*, the family of the genus *Thermolongumtoga*).

The description is the same as for the genus *Thermolongumtoga*. Type genus is *Thermolongumtoga*.

### Description of *Thermolongumtogales* ord. nov.

*Thermolongumtogales* (Ther.mo.lon.gum.to.ga’les. N.L. neut. n. *Thermolongumtoga* type genus of the order; suff.-ales, ending to denote an order; N.L. fem. pl. n. *Thermolongumtogales* the order of the genus *Thermolongumtoga*).

The description is the same as for the genus *Thermolongumtoga*. Type genus is *Thermolongumtoga*.

### Description of *Thermolongumtogae* classis nov

*Thermolongumtogae* (Ther.mo.lon.gum.to’gae. N.L. neut. n. *Thermolongumtoga* type genus of the type order of the class; suff.-ae, ending to denote a class; N.L. fem. pl. n. Thermolongumtogae the class of the order *Thermolongumtogales*).

The description is the same as for the genus *Thermolongumtoga*. Type order is *Thermolongumtogales*.

### Description of *Thermolongumtogota* phyl. nov.

*Thermolongumtogota* (Ther.mo.lon.gum.to.go’ta. N.L. neut. pl. n. *Thermolongumtogae* type class of the phylum; N.L. neut. pl. n. Thermolongumtogota the phylum of the class *Thermolongumtogae*).

The phylum *Thermolongumtogota* is defined based on phylogenetic and phylogenomic analyses of the sole isolated strain J31^T^ and uncultured representatives from various environments. Type order is *Thermolongumtogales*.

## Methods

### Sample collection and “hydrocarbon utilization-driven” strategy-guided isolation

Strain J31 was isolated from venting fluids of Lost City hydrothermal field (30.12°N, 42.12° W). Samples were obtained using the Jiaolong Human-Operated Vehicle on March 17, 2024, and immediately inoculated into a sterile liquid medium, referred to as OM, supplemented with crude oil. Enrichment cultures were incubated at 65 °C. The OM medium contained 0.1 g/L yeast extract, 0.5 g/L tryptone, 0.7 g/L cysteine hydrochloride, and 1 mg/mL resazurin solution with naturally sourced and filtered seawater. The medium was purged with an N2/CO2/H2 gas mixture (80:10:10, v/v), and the pH was adjusted to 7.0 at 25 °C. The culture bottles were sealed with butyl rubber stoppers and screw caps.

After cultured at 65 °C for one month, an initial enrichment was obtained in the OM medium supplemented with crude oil. Based on 16S rRNA gene PCR amplification, the enrichment culture was subsequently transferred into the fresh OM by the limiting dilution method. After an additional month of incubation, a single-cell clone culture was obtained. 3-5 consecutive transfers of this culture under optimal growth conditions (OGM medium, containing 2 g/L maltose, 0.1 g/L yeast extract, 0.5 g/L tryptone, 0.7 g/L cysteine hydrochloride, and 1 mg/mL resazurin solution with naturally sourced and filtered seawater, pH 7.0) were then performed. Finally, the isolation of a pure culture was confirmed by PCR-based identification and electron microscopic observation.

### Phylogenetic analyses

The final, balanced dataset used in this study comprised 226 genomes (206 bacterial references, 15 *Ca.* Bipolaricaulota, and 5 archaeal outgroups) (Tables S6-7, selection is indicated by ‘Yes’ or ‘Representative’). Gene prediction and annotation were performed on all genomes using Prokka (v1.14.6) with the --kingdom Bacteria or --kingdom Archaea flags. OrthoFinder (v2.5.4) was used to identify orthologous protein families (orthogroups, OGs) across the dataset. From this output, we selected 33 OGs that were present in at least 200 of the 226 selected genomes (∼88.5%) (Table S8). Each OG was functionally annotated using eggnog-mapper (v2.1.9). For each protein family, sequences were aligned using MAFFT (v7.487) with the --maxiterate 1000 --genafpair --ep 0 options. Poorly aligned regions were removed from each multiple sequence alignment using trimAl (v1.4.rev15) with the -gappyout option. The trimmed alignments were concatenated into a single supermatrix. A maximum-likelihood phylogenomic tree was inferred from this supermatrix using IQ-TREE (v3.0.0). The best-fit partitioning scheme and substitution models were determined using ModelFinder Plus combined with a merging algorithm to reduce over-parameterization (-m MFP+MERGE). To reduce the computational burden of model selection, a relaxed hierarchical clustering algorithm (-rcluster 10) was used to group data partitions with similar characteristics. Branch support was assessed with 1,000 ultrafast bootstrap replicates (-bb 1000) with an optimization step for each replicate (-bnni). The final tree was visualized using the Interactive Tree of Life (iTOL, v7).

### Microscopic analyses

Cell morphology and structure were observed using phase-contrast microscopy (TS100; NIKON, Japan), confocal laser scanning microscopy (LSM) (LSM900; ZEISS, Germany), field emission scanning electron microscopy (FE-SEM) (Gemini500; ZEISS, Germany), transmission electron microscopy (TEM) (H-7700; Hitachi, Japan) and cryo-electron microscopy (cryo-EM) (Tundra; Thermo Fisher Scientific, USA). Cells in the exponential growth phase in pure culture condition were used for all microscopic observation.

(i) For fluorescence observation, strain J31 was washed with phosphate-buffered saline (PBS) before staining. Cell membranes were stained with DiI at a final concentration of 25 µM, and DNA was stained with DAPI at 10 µg/mL. To track hydrocarbon uptake, strain J31 was cultured with FITC-labeled hexadecane (C16-FITC) at 67 mg/L. (ii) For SEM observation, strain J31 was fixed with 2.5% glutaraldehyde, washed three times with PBS, gradually dehydrated in ethanol series (30%, 50%, 70%, 90%, and 100% for 10 min at each step), and subsequently imaged. (iii) For TEM observation, strain J31 was fixed with 2.5% glutaraldehyde in 0.1 M sodium cacodylate buffer (pH7.4) at 4 °C for 12 h, followed by post-fixation with 1% osmium tetroxide and potassium hexacyanoferrate in cacodylate buffer. Cells were then dehydrated in an alcohol series and embedded into Epon. Ultrathin sections were collected on copper grids, stained with uranyl acetate and lead citrate, and visualized. (iv) For cryo-EM observation, 2 μL of cell culture were placed on glow-discharged 300-mesh Au grid (Quantifoil R1.2/1.3) in a Vitrobot Mark IV (Thermo Fisher Scientific, USA). Vitrification was performed at 25 °C under 100% humidity. Samples were blotted for 2.0 s and plunge-frozen into liquid ethane. The frozen Au grids were mounted onto a liquid-nitrogen cryo-specimen holder and loaded into a Tundra microscope equipped with a cold-field emission electron gun operating at 100 kV.

### Quantitative PCR

SYBR green-based real-time PCR was performed on the QuantStudio^TM^ 6 Flex (Thermo Fisher Scientific, USA) using the SYBR^®^Green Realtime PCR Master Mix (Toyobo, Japan) to quantify the population of strain J31. The forward and reverse primers, J31F (5′-CCGTTACCCCACCAACTAGC-3′) and J31R (5′-CGGGCTAATA CCCGATGGTC-3′), were designed and synthesized based on the 16S rRNA gene sequences of strain J31. Total DNA was extracted from pure cultures using the conventional phenol extraction method. Standard curves for quantification were determined based on 10-fold serial dilutions of the target PCR products of strain J31 at known concentrations. Cell growth was estimated using 16S rRNA gene copy number as a proxy for cell population. All analyses were conducted in triplicate.

### Growth characteristics

Various substrates were added to the basal OM medium. The following single carbon sources, all at a final concentration of 0.2% (w/v), were tested: maltose, lactose, trehalose, galactose, gelatin, pectin, cellulose, laminarin, fucoidan, xylan, glucose, rhamnose, xylose, and arabinose. 20 mM of sodium sulfate, sodium acetate and sodium nitrate were individually added to the OM medium and tested. Sensitivities to kanamycin (50 µg/mL), gentamicin (50 µg/mL), chloramphenicol (25 µg/mL), hygromycin (50 µg/mL), ampicillin (100 µg/mL), rifampin (50 µg/mL), lovastatin (5 µg/mL), and simvastatin (5 µg/mL) were determined in the OGM medium. The effects of temperature, pH, and concentrations of NaCl on cell growth of strain J31 were also measured in the OGM medium. Cell growth was estimated using 16S rRNA gene copy number by quantitative PCR. All analyses were conducted in triplicate.

### Genome sequencing and analysis

Genomic DNA of strain J31 was extracted using Bacterial/fungal DNA extraction kit (Majorbio, China) according to manufacturer’s protocol. Purified genomic DNA was sequenced using a combination of PacBio Sequel IIe and Illumina sequencing platforms. Sequence assembly was performed with GUnicycle v0.4.8. Gene prediction and annotation were carried out using Glimmer or Prodigal v2.6.3. tRNA-scan-SE (v2.0) was used for tRNA gene prediction, and Barrnap (v0.9) was used for rRNA gene prediction. Protein-coding sequences (CDSs) were annotated by alignment against the NR, Swiss-Prot, Pfam, GO, COG, KEGG and CAZy databases using sequence alignment tools such as BLAST, Diamond and HMMER. Briefly, each set of query proteins were aligned with the databases, and annotations from the best-matched subjects (E-value < 10^-5^) were assigned to the genes. Biosynthetic gene clusters (BGCs) of secondary metabolites were identified by antiSMASH v5.1.2 software.

### *n*-Alkanes extraction and analysis

Throughout the entire sample processing, all glassware used was combusted at 400 °C for 4 h to remove any trace of organic matter (*40*). Serpentine minerals were firstly cleaned with *n*-hexane, and then dried at 70 °C for 12 h to remove surface contaminants. Hydrocarbons from serpentine minerals were extracted with *n*-hexane for 72 h, concentrated using a vacuum rotary evaporator, and subsequently analyzed by gas chromatography-mass spectrometry (GC-MS) (7890A-5975C, Agilent, USA). All extractions and subsequent quantifications were performed in triplicate. The GC temperature program was as follows: injection at 80 °C followed by 2 min of constant temperature, gradual temperature change from 80 °C to 290 °C at 4 °C per min, and then another 20 min of constant temperature. The MS system was an EI model operated at 70 eV.

### Colloidal gold immunoelectron microscopy

Strain J31 in OM medium was incubated with FITC or FITC-labeled hexadecane (C16-FITC) at the final concentration of 67 mg/L for 4 h at 65 °C. Cells were subjected to collection, fixation, and embedding, followed by preparation of ultrathin sections. The sections were incubated with the primary antibody (anti-FITC monoclonal antibody, MA5-14696, Thermo Fisher Scientific, USA), and unbound antibody was removed by washing. Subsequently, the sections were incubated with a colloidal gold-labeled secondary antibody that specifically binds the primary antibody. After washing away unbound secondary antibody, the sections were stained with uranyl acetate and observed by TEM. Localization of the target antigen (FITC or C16-FITC) was visualized by the presence of colloidal gold particles.

### Transcriptome analysis

Strain J31, incubated with or without hexadecane (67 mg/L) for 4 h, was harvested by centrifugation at 4 °C, and used for transcriptomic analysis by Majorbio (Beijing, China). Briefly, total RNA was extracted using CTAB method, and genomic DNA was removed. RNA-seq library was prepared and sequenced with the Illumina Novaseq 6000 (Illumina Inc., San Diego, CA, USA). The sequencing data generated from Illumina platform were subjected to bioinformatics analysis on Majorbio Cloud Platform (www.majorbio.com). Gene and transcript abundances from RNA-Seq data were quantified using RSEM. The differentially expressed genes were identified using the edgeR, DESeq2, or DESeq packages. All analyses were conducted in triplicate.

### Isotope tracing untargeted metabolomics

Strain J31 was incubated with 67 mg/L hexadecane-1,2-^13^C2 (CAS Number: 158563-27-0, Sigma-Aldrich, USA) or unlabeled hexadecane for 2 weeks. Bacterial pellets (about 10^7^ bacteria) were collected by centrifugation at 4 °C, quickly frozen in liquid nitrogen, and subsequently subjected to metabolomic analysis by Shanghai Biotree Biotech (Shanghai, China). Liquid chromatography–tandem mass spectrometry (LC–MS/MS) analyses were performed using an UHPLC system (Vanquish, Thermo Fisher Scientific, USA) with a Phenomenex Kinetex C18 (2.1 mm × 100 mm, 2.6 μm) coupled to Orbitrap Exploris 120 mass spectrometer (Orbitrap MS, Thermo). The raw data were converted to the mzXML format using ProteoWizard and processed with an in-house R package. Metabolite identification was conducted against the in-house MS2 database (Biotree DB). All analyses were conducted in quadruplicate.

### Annotation of hydrocarbon-degrading genes

Genes involved in hydrocarbon degradation within the genomes or metagenome-assembled genomes (MAGs) were identified by searching for sequence homology against the CANT-HYD database, as proposed by Khot and colleagues (*47*) and publicly available in their GitHub repository (https://github.com/dgittins/CANT-HYD-HydrocarbonBiodegradation). Homology searches were performed for each genome or MAG using the *hmmsearch* function with an E-value cutoff of 1×10^10^.

### Co-cultivation of strain J31 with serpentine minerals

Serpentine minerals used in this study were collected from the Mussau Trench in the Western Pacific (2.42° N, 148.76° E) and identified as serpentinite based on cross-polarized microscopic observations and elemental composition analyses. Strain J31 was cultured in OM medium supplemented with 25 g/L (w/v) serpentine minerals for 4 weeks. Colonization of strain J31 on the serpentine surfaces was examined by SEM. Black precipitates in the culture supernatants were collected by centrifugation, fried, and analyzed by energy dispersive spectroscopy (EDS, Gemini500, ZEISS, Germany). For elemental analysis, culture supernatants were collected by centrifugation and treated with HNO₃ to remove organic matter. The concentrations of soluble iron, silicon, and magnesium were then determined by the inductively coupled plasma optical emission spectroscopy (ICP-OES, UlltiMate3000-iCAP TQ, Thermo Fisher Scientific, USA). All analyses were conducted in triplicate.

### Isolation and identification of the active compound produced by strain J31

Strain J31 was cultured in the 2 L anaerobic bottle containing 2 g/L maltose, 0.1 g/L yeast extract, and 0.5 g/L tryptone in naturally sourced and filtered seawater (pH 7.0) at 65 °C for 10 days. The culture supernatant was collected by centrifugation and extracted with ethyl acetate. After filtration and removal of the solvent using a rotary evaporator under vacuum at 45 °C, the crude extract was fractionated using a methanol–water gradient by a reversed-phase high-performance liquid chromatography (RP-HPLC) system (Agilent, USA) with an Eclipse XDB-C18 column (5 μm, 9.4 × 250 mm; Agilent, USA). The elution fractions were collected, and tested for their antimicrobial activity against *Ca.* Bipolaricaulota strain GYR-2 and *Thermotoga* strain GYR-1, both of which were previously isolated from deep-sea hydrothermal fields by our team.

### Data availability

The genome sequencing data were deposited in the NCBI database (PRJNA1344826). The RNA sequencing data have been deposited in the NCBI database (PRJNA1344839). The metabolomics data have been deposited to MetaboLights repository with the study identifier MTBLS13250.

## Acknowledgements

This work was supported by the NSFC Innovative Group Grant (No. 42221005), Science and Technology Innovation Project of Laoshan Laboratory (Grant Nos. LSKJ202203103 and 2022QNLM030004-3), Shandong Provincial Natural Science Foundation (Grant No. ZR2024ZD49), Major Research Plan of the National Natural Science Foundation (Grant No. 92351301), Qingdao West Coast New District University Presidents Fund (Grant No. E42424101N), Taishan Scholars Program (Grant No. tstp20230637) and the NSFC Grant (No. 42406128).

## Author contributions

G.L., J.Z., S.M., and C.S. conceived the study and wrote the manuscript. G.L. and J.Z. carried out most of cultivation and culture-based experiments. Y.S. performed the phylogenetic analyses. R.L. and Y.G. conducted the bioinformatics analyses. S.Y. conducted sampling and identification of serpentinite. H.W., M.L., and H.Q. carried out the venting fluids sampling. Q.P. conducted GC-MS analysis. All authors have read and approved the manuscript submission.

## Competing interests

The authors declare no competing interests.

---

**Fig. S1.**
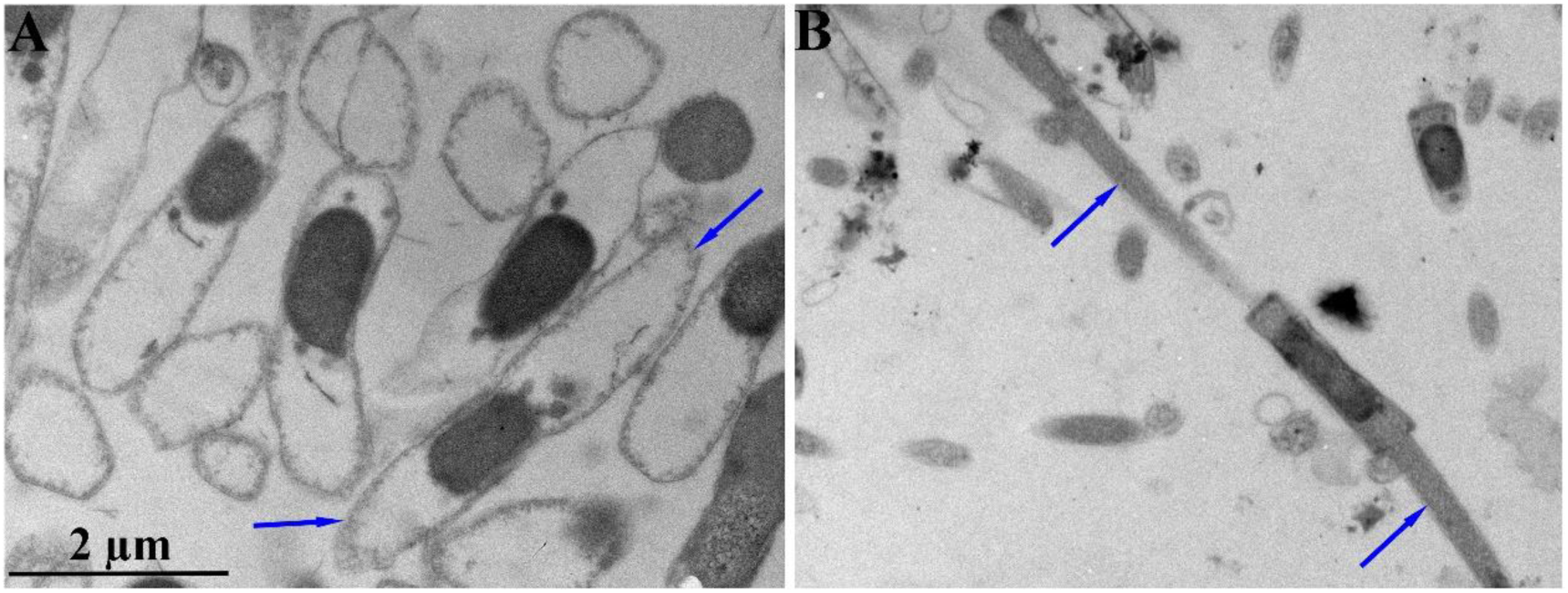
Morphological similarity between *Thermotogota* and *Ca.* Bipolaricaulota members. (**A** and **B**) Transmission electron micrographs of ultrathin sections of *Thermotogales* sp. GYR-1 (A) and strain J31 (B). The blue arrows indicate the Toga structures.

**Fig. S2.**
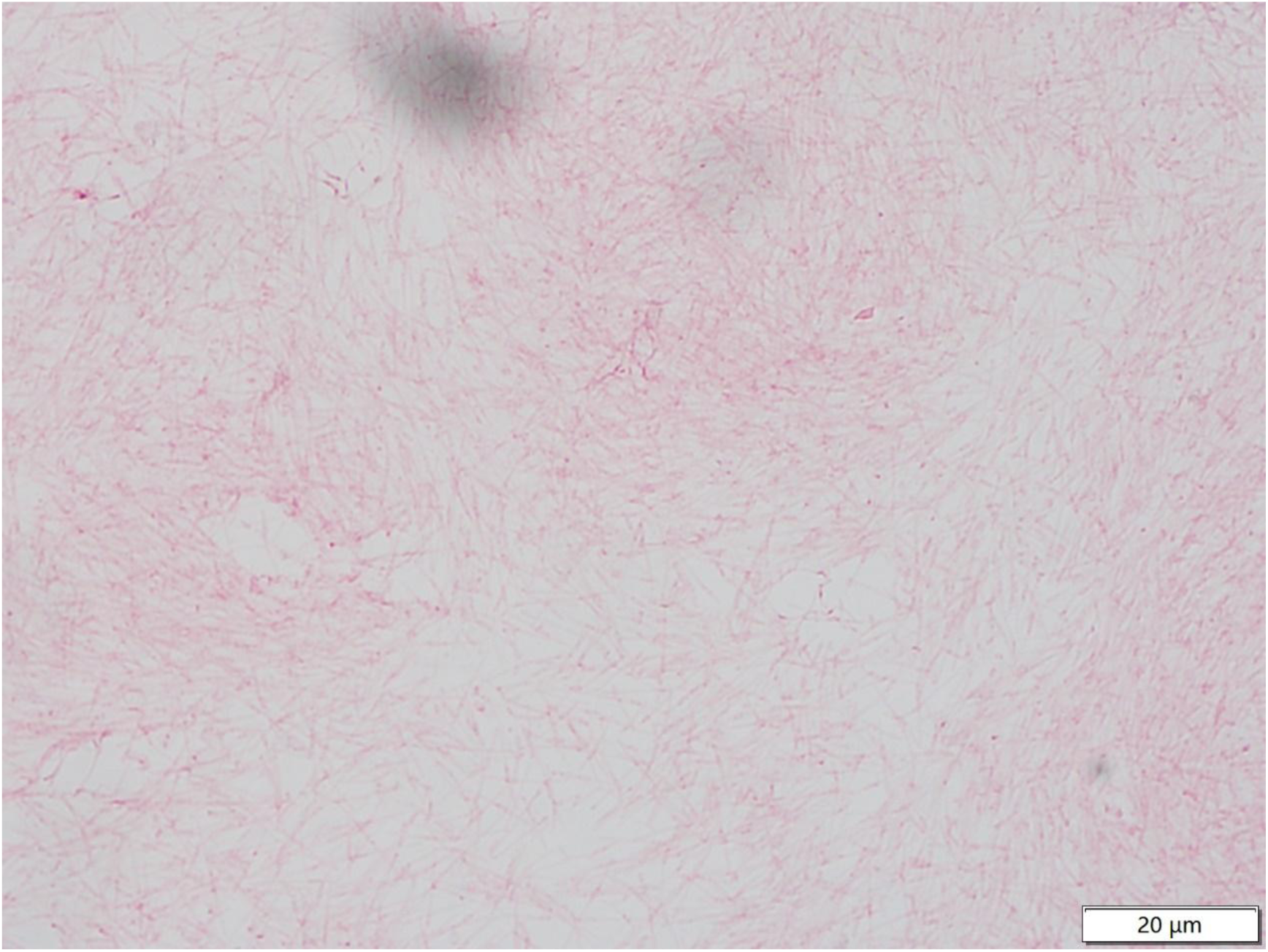
Micrographs showing strain J31 is Gram-negative. Stained cells on the glass slide were observed under an optical microscope using the oil immersion objective (100 ×).

**Fig. S3.**
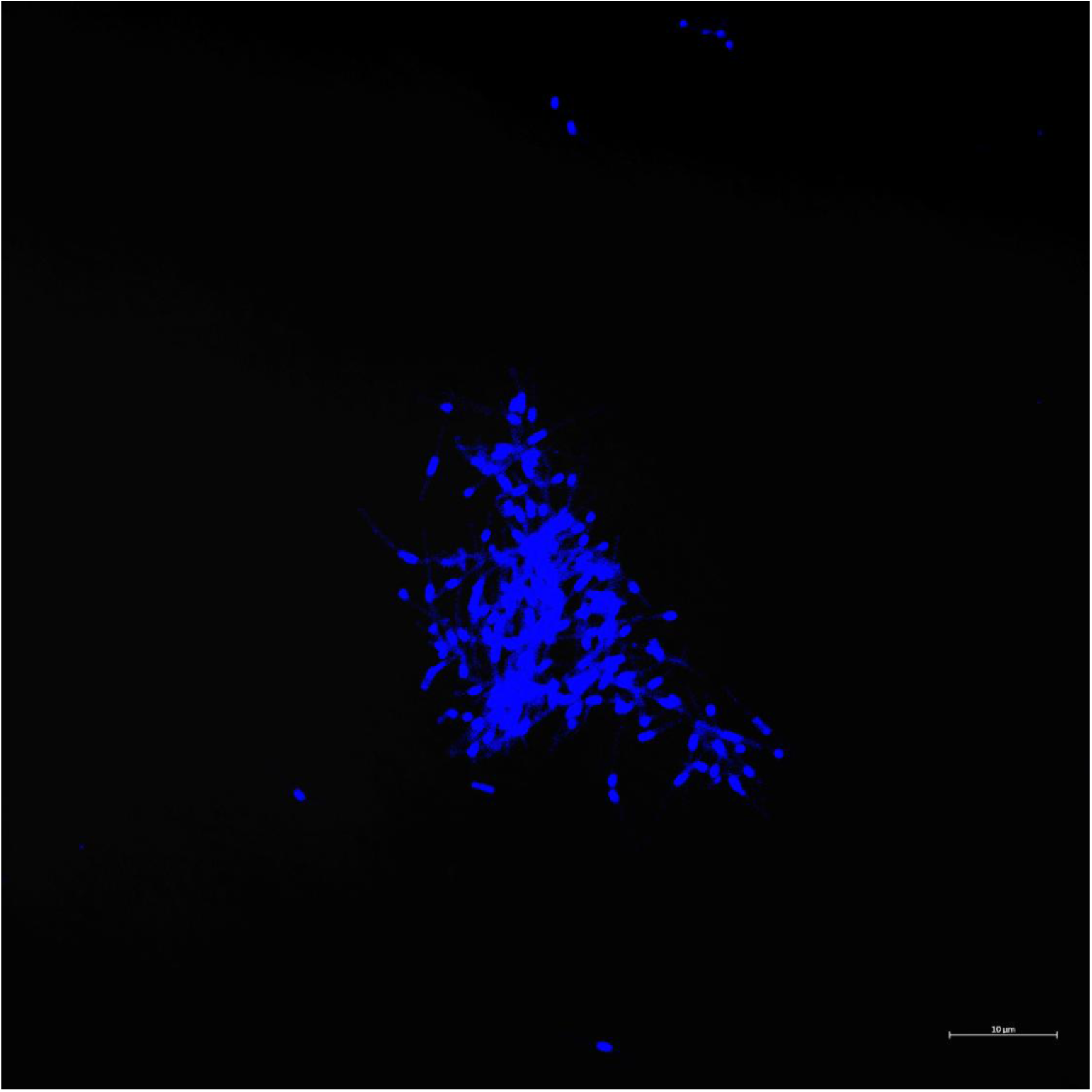
Fluorescence micrographs showing the localization of nucleic acids in the central rod of strain J31. Nucleic acids were stained with DAPI and visualized by LSM.

**Fig. S4.**
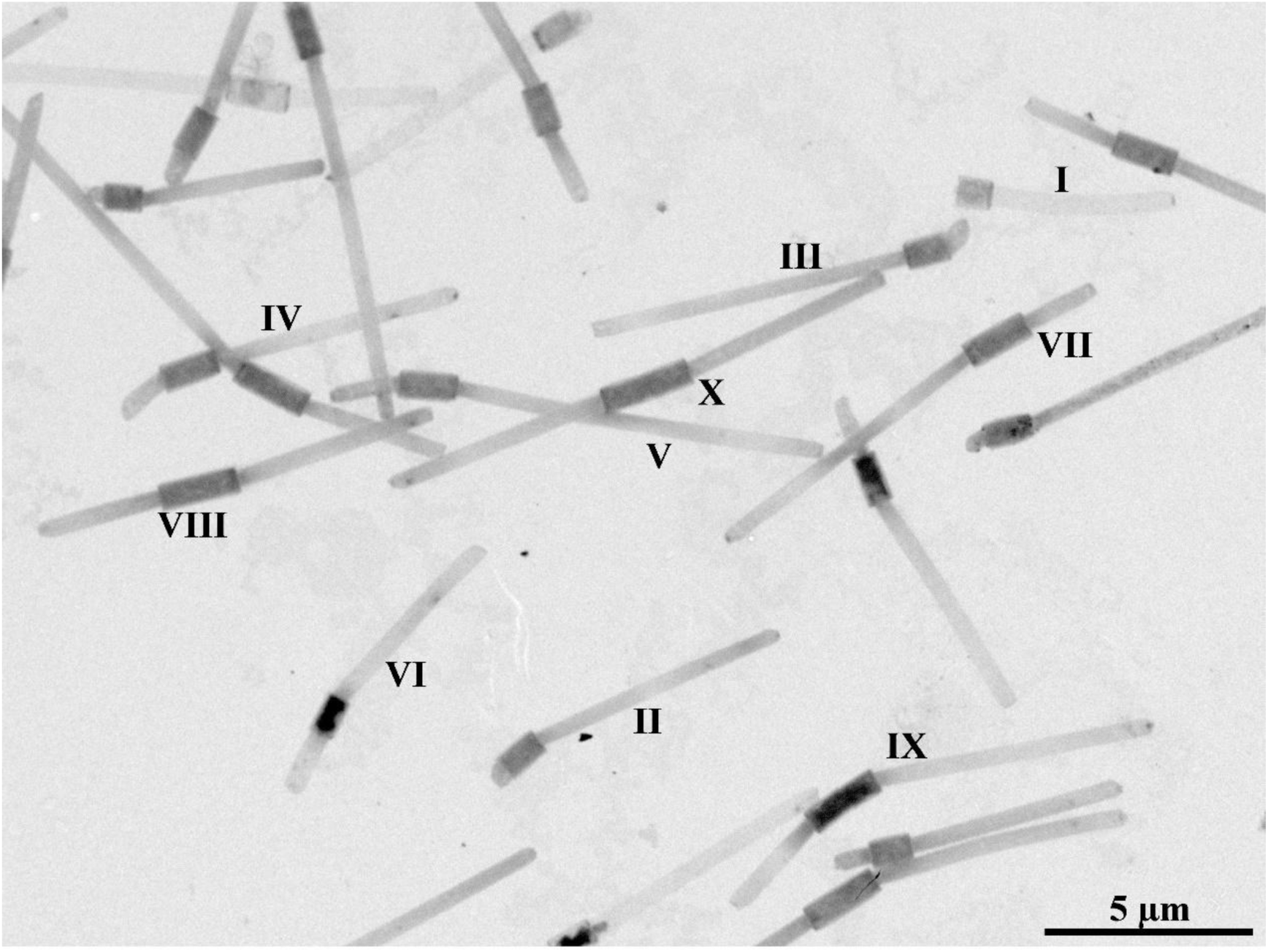
TEM images showing different morphologies of strain J31 at different growth stages. Shapes I–X show the division process of strain J31.

**Fig. S5.**
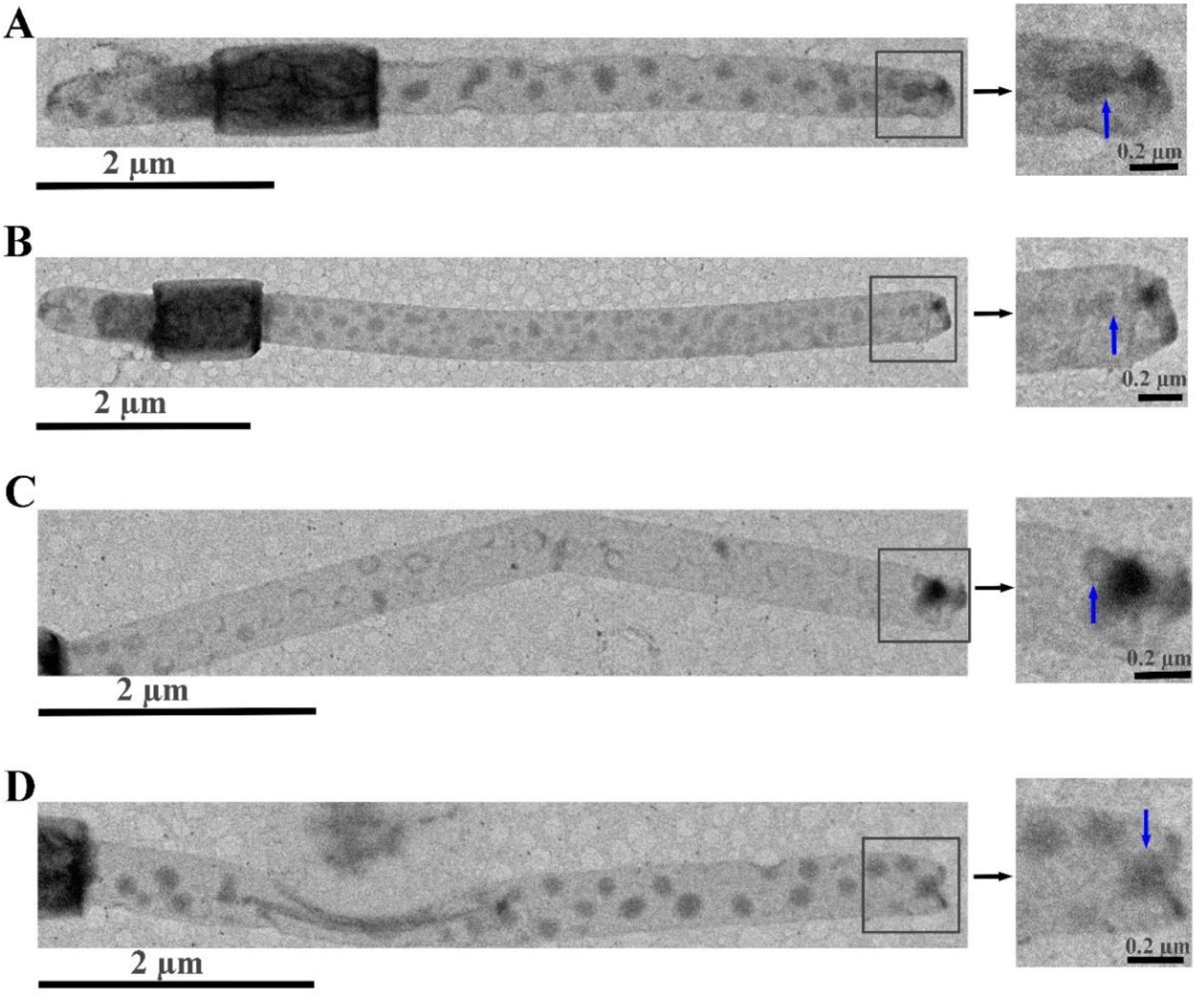
TEM images showing vesicle-like structures of crude oil budding from Toga ends in strain J31. (**A** to **D**) Different J31 cells with vesicle-like structures of crude oil in Toga observed by TEM. Enlarged views of the boxed regions were shown in graph on the right. The blue arrow indicates the vesicle formation process.

**Fig. S6.**
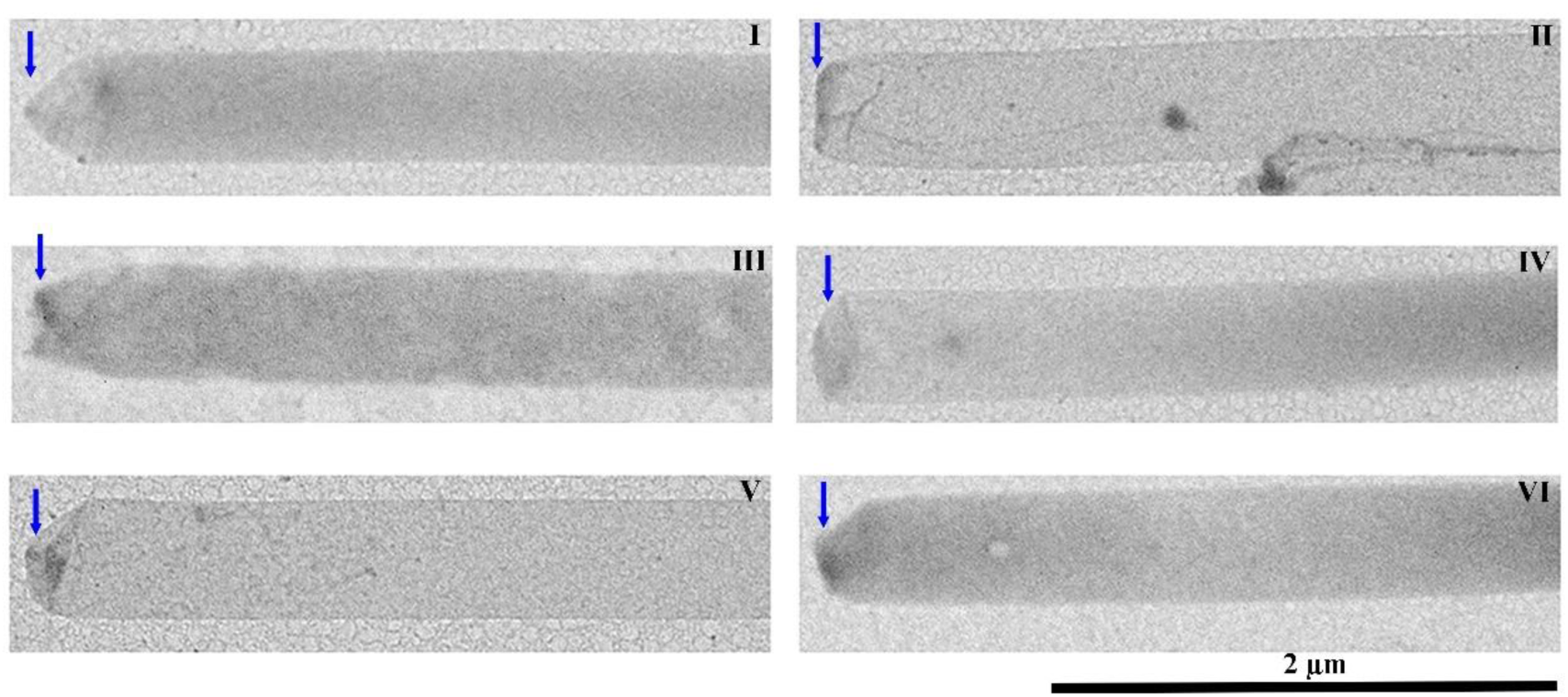
TEM images showing the different morphologies of Toga ends in strain J31. Panels I-VI display the morphological progression of the Toga ends during external nutrient absorption, with blue arrows indicating the sequence of these changes.

**Fig. S7.**
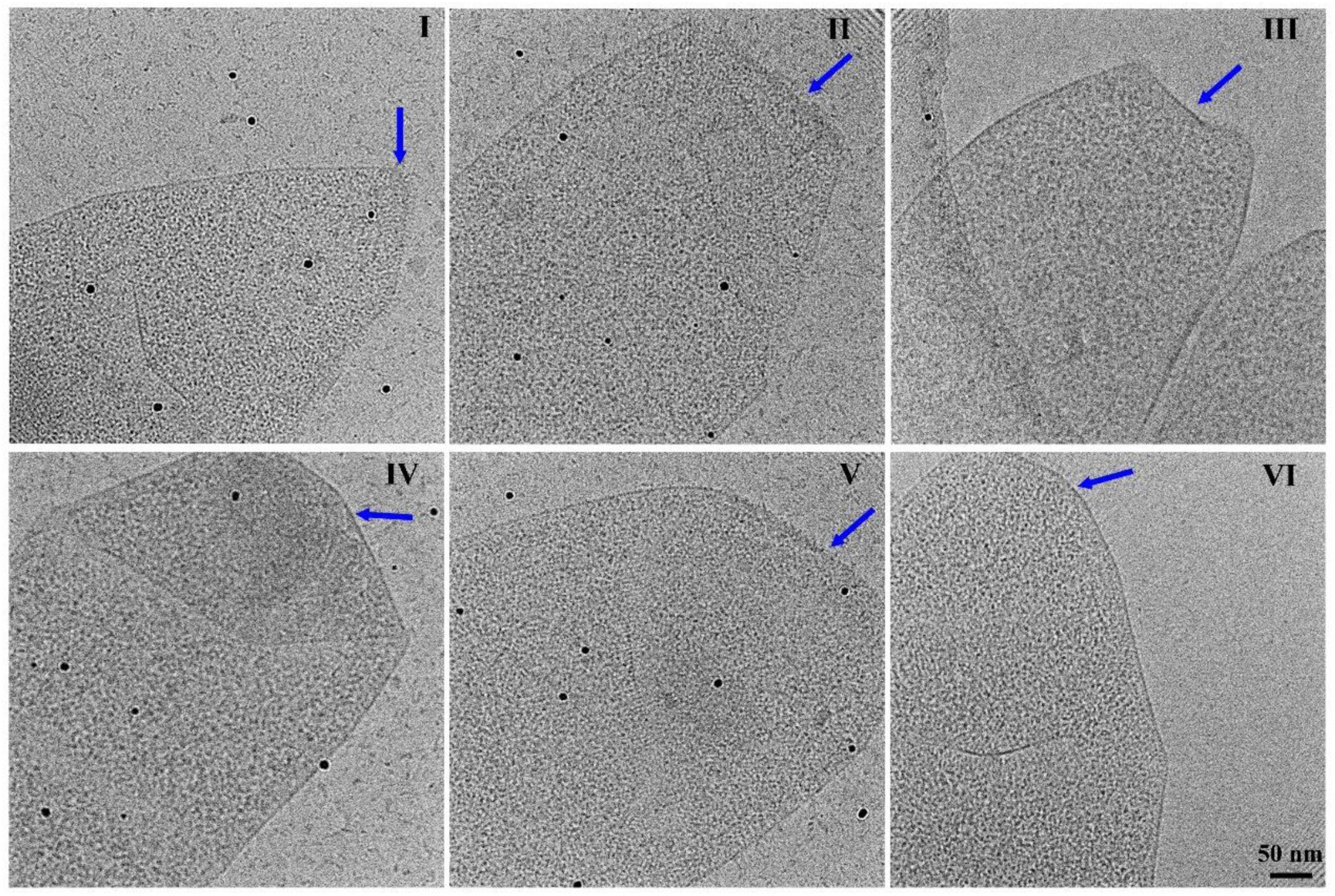
Cryo-EM revealing the different morphologies of Toga ends in strain J31. Panels I-VI show the morphological progression of Toga ends during external nutrient absorption, with blue arrows indicating the sequence of these changes.

**Fig. S8.**
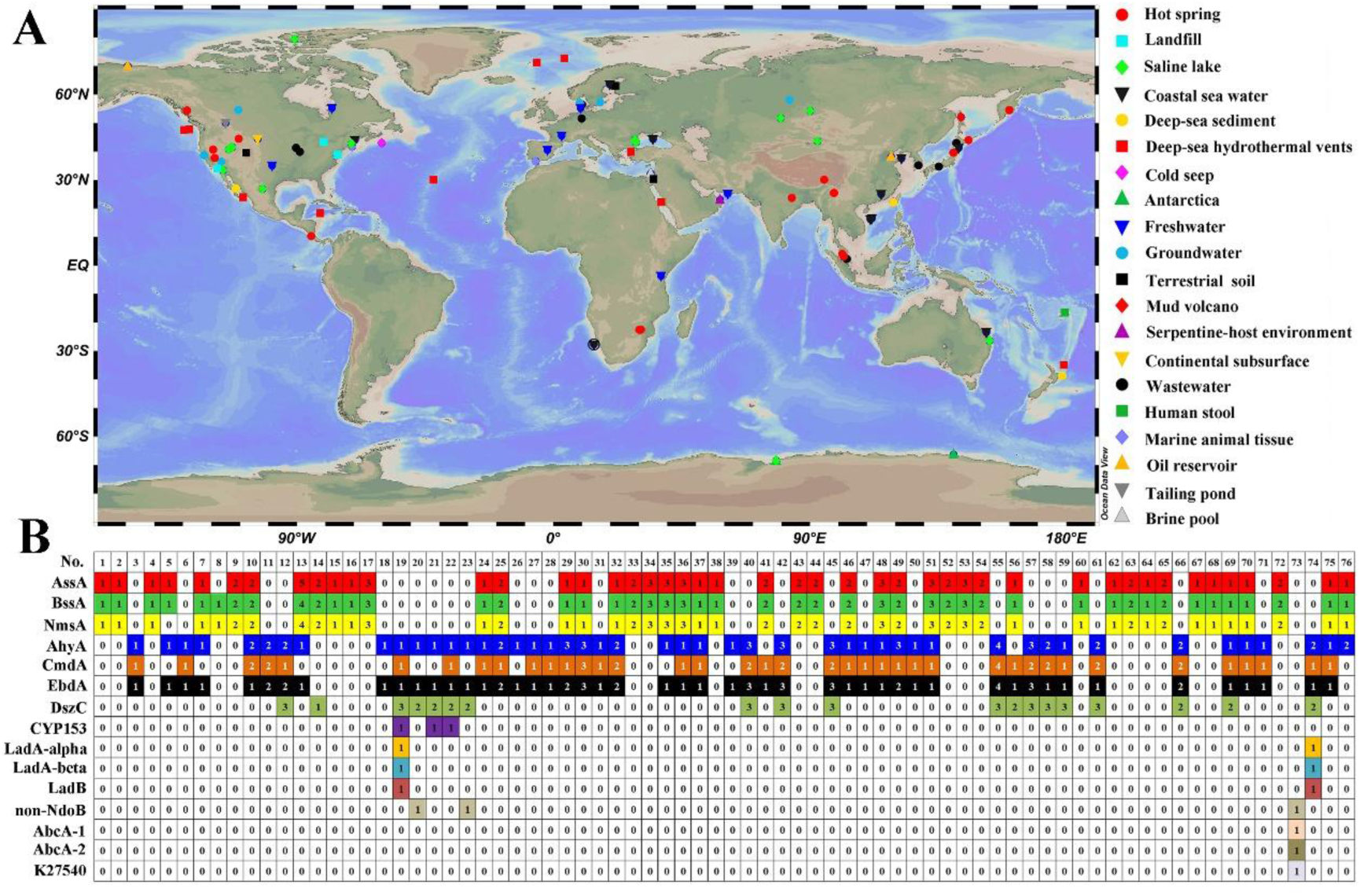
Global distribution and hydrocarbon utilization potential analysis of *Ca.* Bipolaricaulota. (**A**) Ninety-nine 16S rRNA gene sequences representing *Ca.* Bipolaricaulota were obtained from the NCBI database (Table S2) and their geographic locations were summarized. The map was generated by Ocean Data View (Schlitzer, Reiner, Ocean Data View, https://odv.awi.de, 2025). (**B**) A total of 189 complete and incomplete genomes (MAGs) of *Ca.* Bipolaricaulota from NCBI database were obtained (Table S3), and they were analyzed using the CANT-HYD database (https://github.com/dgittins/CANT-HYD-HydrocarbonBiodegradation). The numbers in the colored box represent gene counts. Abbreviations: AssA, alkyl-succinate synthase; BssA, benzyl-succinate synthase; NmsA, naphtylmethyl succinate synthase; AhyA, molybdopterin-family alkane C2 methylene hydroxylase; CmdA, molybdopterin-family ethylbenzene dehydrogenase subunit alpha; EbdA, molybdopterin-family ethylbenzene dehydrogenase subunit alpha; DszC, dibenzothiophene monooxygenase; CYP153, alkane oxidizing cytochrome P450; LadA-alpha, long-chain alkane hydrolase A subunit alpha; LadA-beta, long-chain alkane hydrolase A subunit beta; LadB, long-chain alkane hydrolase B; non-NdoB, benzene/toluene/naphthalene dioxygenase subunit alpha; AbcA-1/2, benzene carboxylase; K27540, naphthalene carboxylase.

**Fig. S9.**
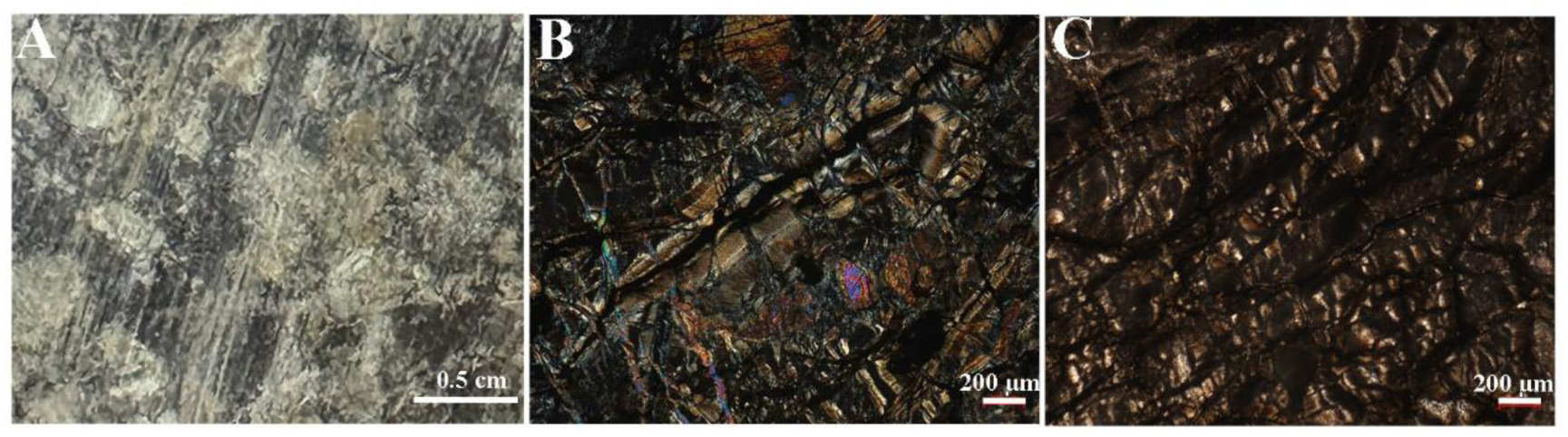
Images of serpentinite samples used in this study. (**A**) Hand specimen of serpentinite, showing clearly visible serpentine minerals. (**B** and **C**) Photomicrographs of thin sections under cross-polarized light, revealing extensive serpentinization of the protolith. The characteristic mesh texture typical of serpentinite is clearly observable in the thin sections.

**Fig. S10.**
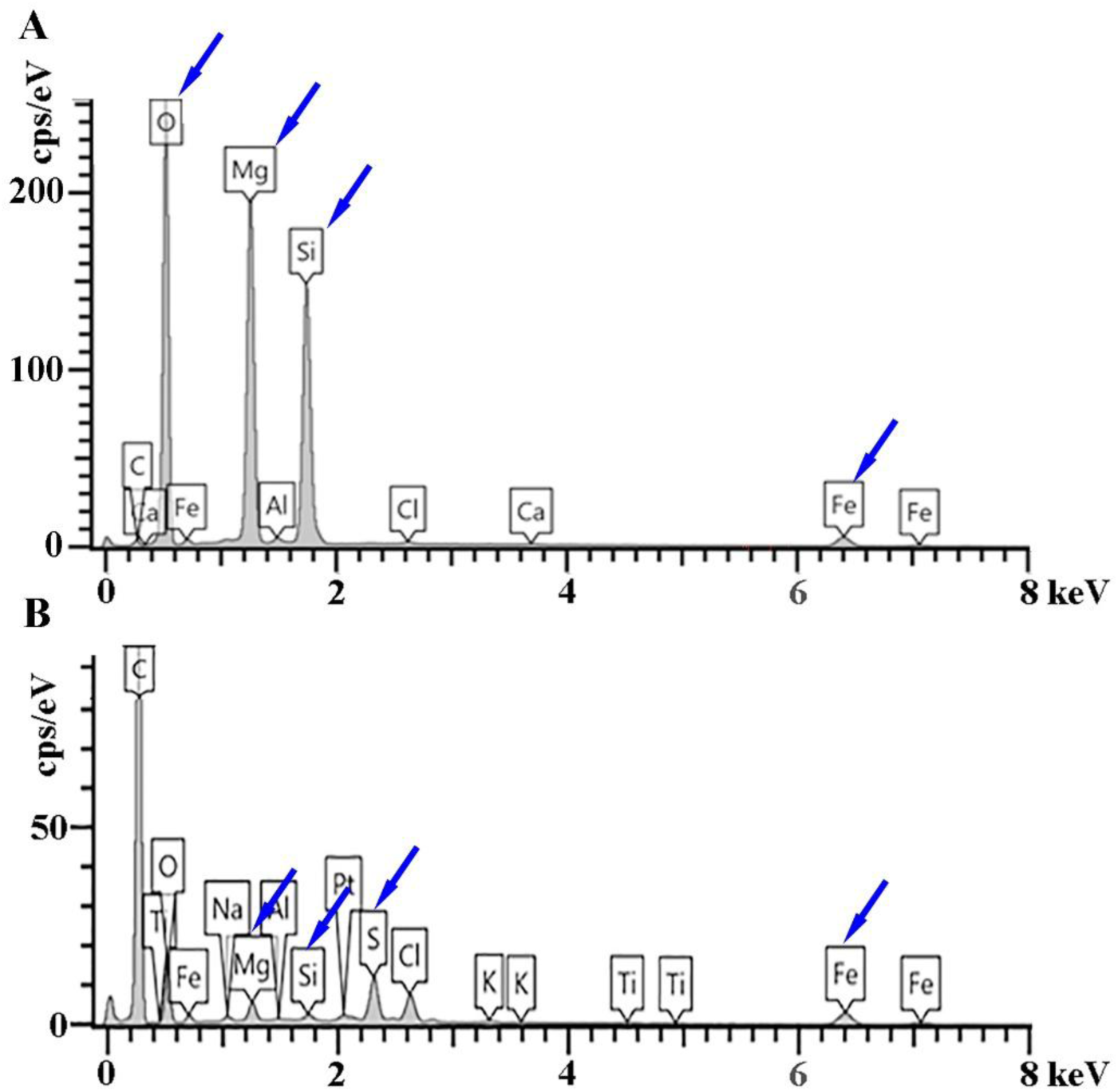
Elemental composition of serpentine minerals and co-culture precipitates. (**A** and **B**) Energy dispersive spectroscopy (EDS) analysis diagrams of serpentine minerals used in the study (A) and black precipitates induced by strain J31 (B). The blue arrows indicate the main elements.

**Fig. S11.**
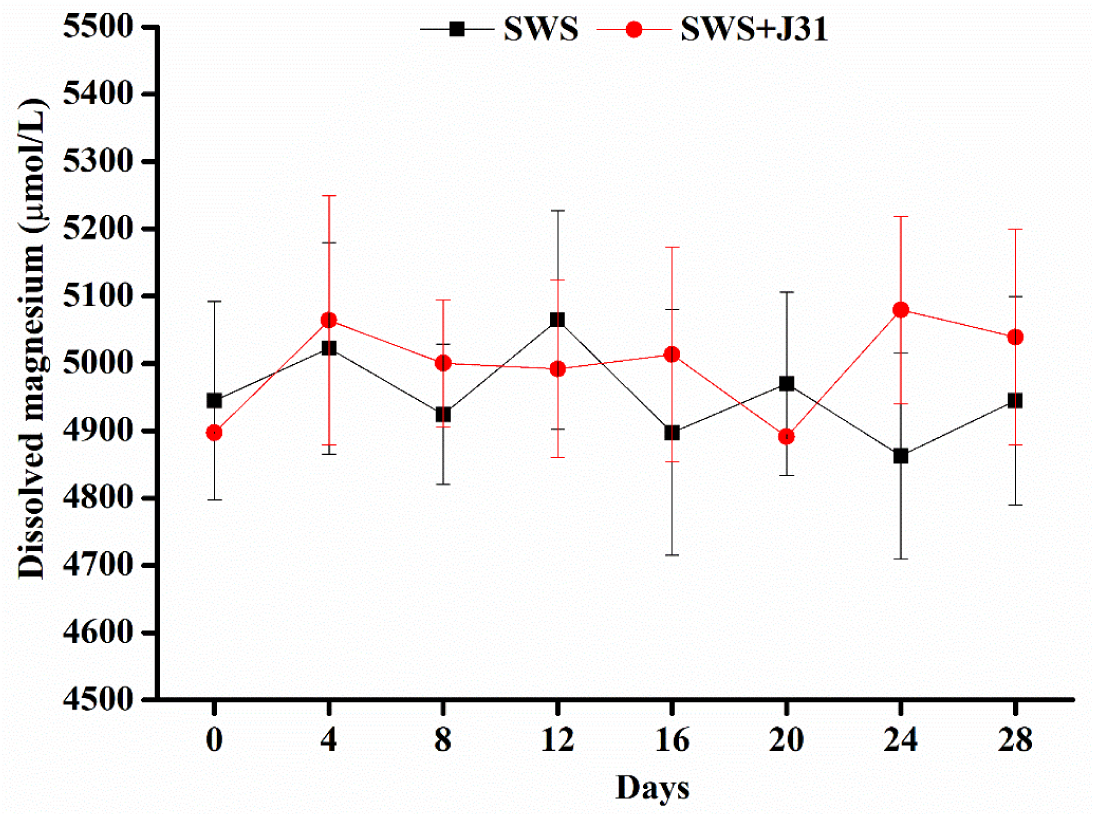
Analysis of magnesium release from serpentine minerals. The magnesium content in the culture supernatants was analyzed by inductively coupled plasma optical emission spectroscopy (ICP-OES). SWS refers to serpentine minerals.

**Fig. S12.**
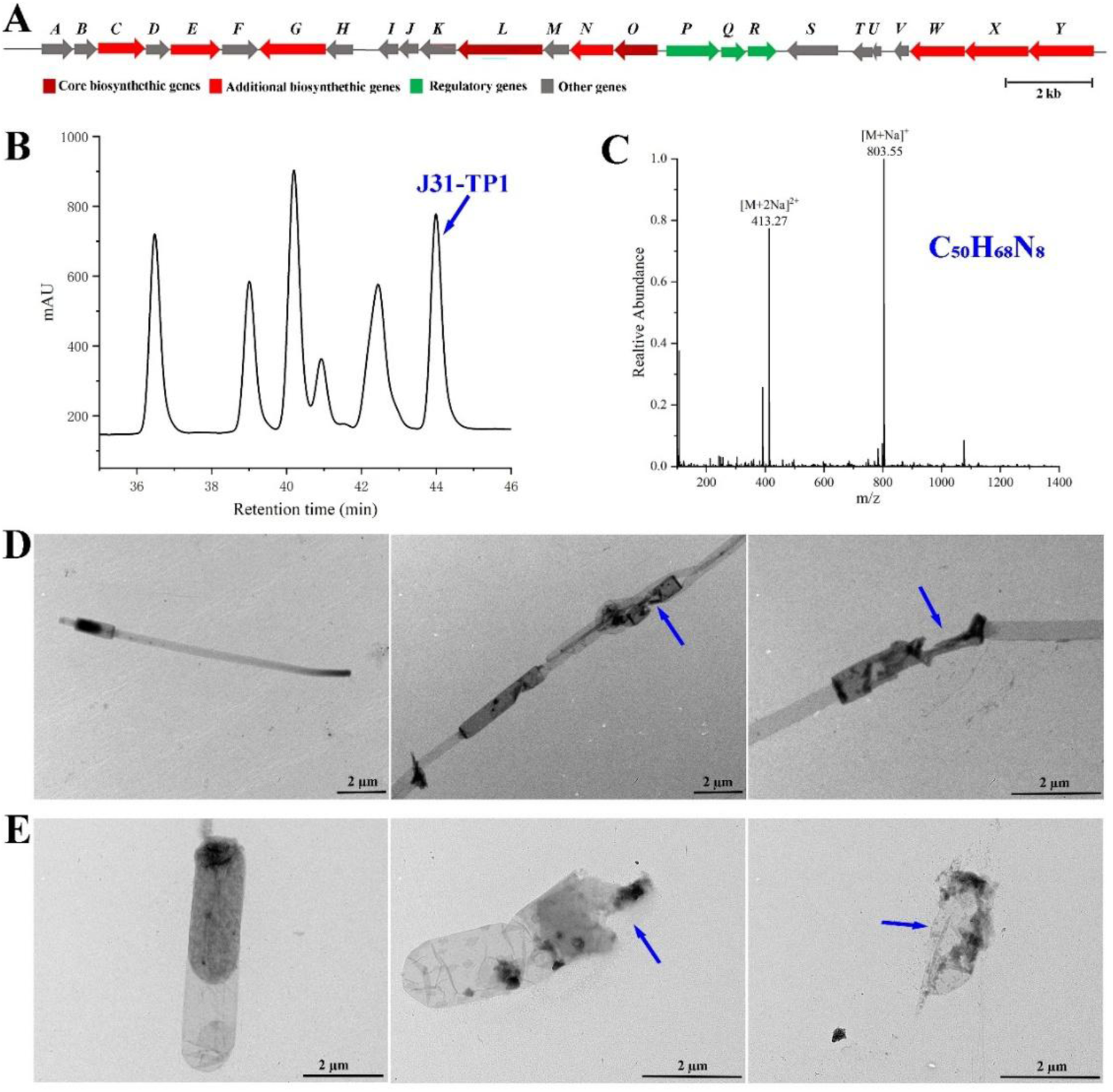
Strain J31 produces an active terpenoid alkaloid-like compound inhibiting the growth of Toga-containing bacteria. (**A**) Complete biosynthetic gene cluster involved in the production of terpenoids in the genome of strain J31. Products of these genes: A, tryptophan synthase alpha chain; B, shikimate kinase; C, hypothetical protein; D, type II 3-dehydroquinatedehydratase; E, aspartate aminotransferase; F, prephenate dehydratase; G, GMP synthase; H, xanthine phosphoribosyl-transferase; I, DUF116 domain-containing protein; J, YjbQ family protein; K, 4-hydroxy-3-methylbut-2-enyl diphosphate reductase; L, squalene-hopene cyclase; M, hypothetical protein; N, adenosyl-hopene transferase; O, polyprenyl synthetase family protein; P, hypothetical protein; Q, response regulator transcription factor; R, hypothetical protein; S, IS110 family transposase; T, PIN domain-containing protein; U, ribbon-helix-helix protein; V, DUF1667 domain-containing protein; W, FAD-dependent oxidoreductase; X, NAD(P)/FAD-dependent oxidoreductase; Y, glycerol kinase. (**B**) Chromatograph profile of the active substance (designated J31-TP1) isolated from strain J31 on a reversed-phase C18 column eluted with a methanol-water gradient at a flow rate of 2.0 mL/min. (**C**) High-resolution mass spectrometry and possible molecular formula (C50H68N8) of J31-TP1. (**D** and **E**) J31-TP1 significantly induced the membrane rupture of another strain belonging to *Ca*. Bipolaricaulota (strain GYR-2, D), and *Thermotoga* strain GYR-1 (E), both of which are thermophilic bacteria inhabiting the serpentinization-influenced environments. Blue arrows indicate the membrane rupture.

**Fig. S13.**
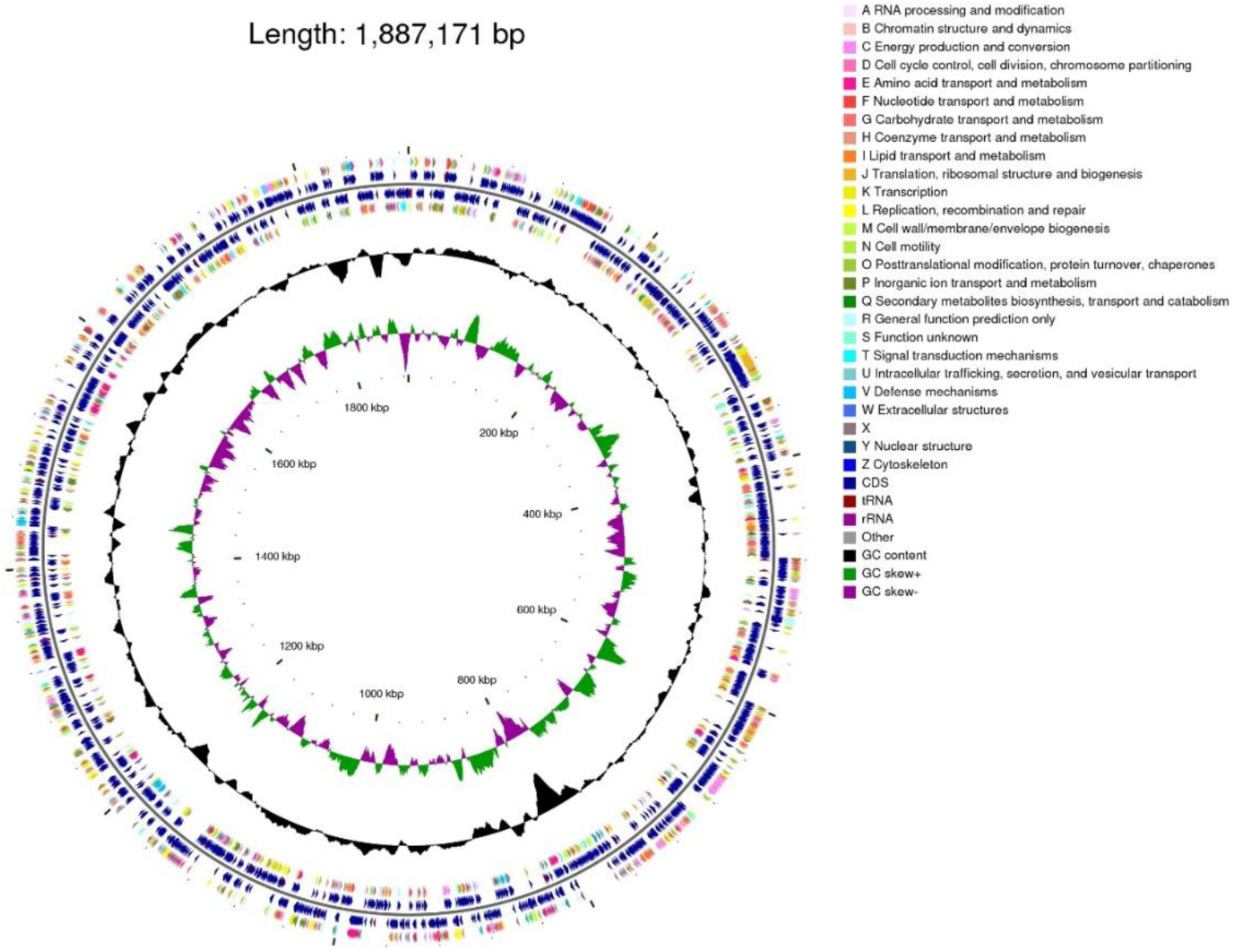
Circular representation of the strain J31 genome. Circles from the center to the outside: 1, genome size; 2, GC skew; 3, GC content; 4, ncRNA, CDS and gene function annotation.

**Table S1.**
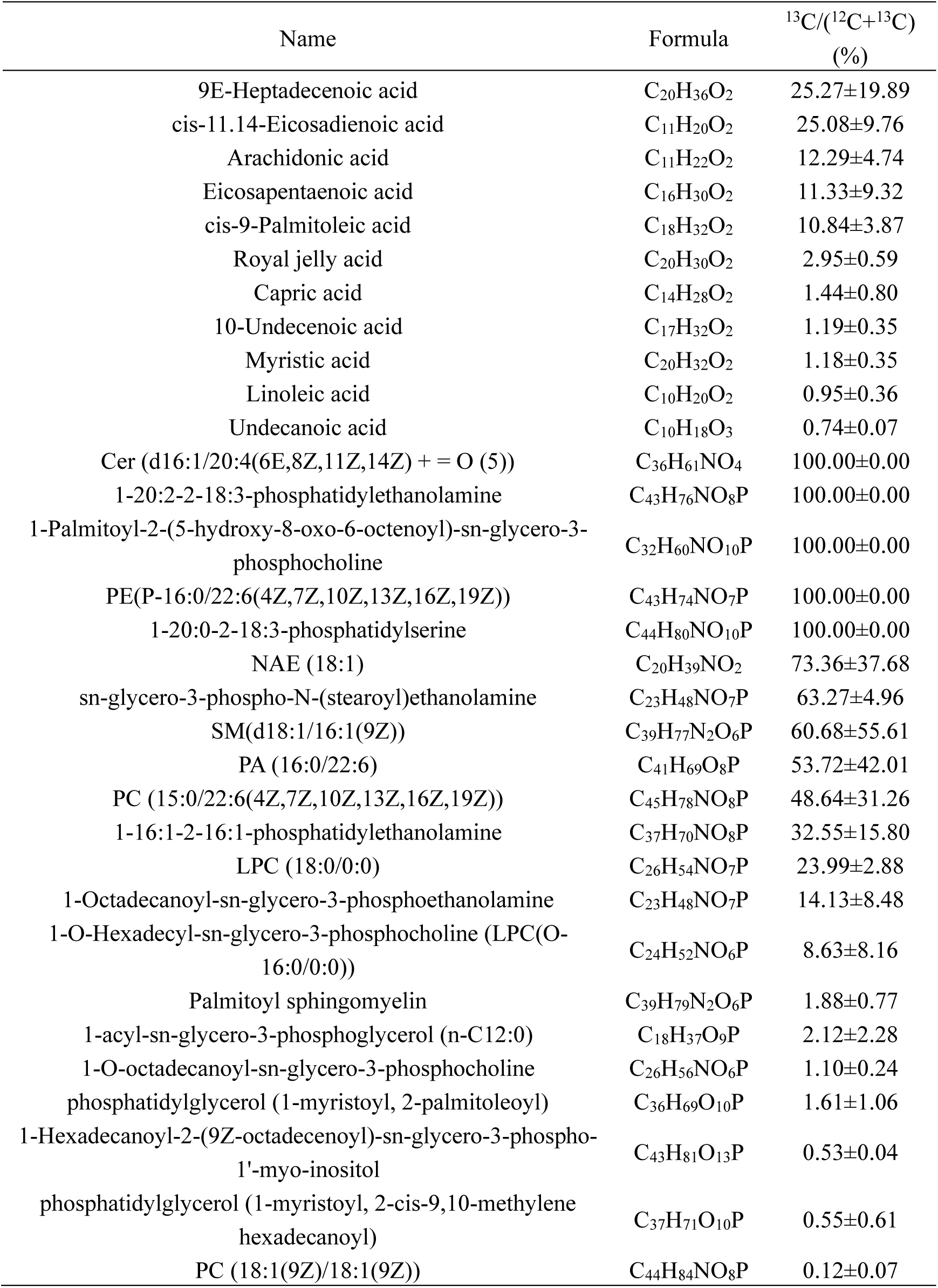
Free fatty acids and lipids labeled with 13C were detected in strain J31 with hexadecane-1,2-^13^C2 as a tracer.

**Table S5.** Composition of *n*-alkanes extracted from serpentine minerals.

| *n*-alkanes | Formula | ng/g serpentine |
| --- | --- | --- |
| C11 | C<sub>11</sub>H<sub>24</sub> | 1.62±0.44 |
| C12 | C<sub>12</sub>H<sub>26</sub> | 6.20±0.85 |
| C13 | C<sub>13</sub>H<sub>28</sub> | 4.16±0.80 |
| C14 | C<sub>14</sub>H<sub>30</sub> | 4.02±0.63 |
| C15 | C<sub>15</sub>H<sub>32</sub> | 1.48±0.21 |
| **C16** | **C<sub>16</sub>H<sub>34</sub>** | **11.70±1.24** |
| C17 | C<sub>17</sub>H<sub>36</sub> | 4.86±0.56 |
| C18 | C<sub>18</sub>H<sub>38</sub> | 4.15±0.32 |
| C19 | C<sub>19</sub>H<sub>40</sub> | 1.27±0.21 |
| C20 | C<sub>20</sub>H<sub>42</sub> | 2.61±0.12 |
| C21 | C<sub>21</sub>H<sub>44</sub> | 0.63±0.00 |
| C22 | C<sub>22</sub>H<sub>46</sub> | 0.71±0.12 |
| C23 | C<sub>23</sub>H<sub>48</sub> | 1.13±0.12 |
| C24 | C<sub>24</sub>H<sub>50</sub> | 0.56±0.12 |
| C25 | C<sub>25</sub>H<sub>52</sub> | 0.78±0.24 |
| C26 | C<sub>26</sub>H<sub>54</sub> | 0.92±0.12 |
| C27 | C<sub>27</sub>H<sub>56</sub> | 0.99±0.24 |
| C28 | C<sub>28</sub>H<sub>58</sub> | 0.71±0.12 |
| C29 | C<sub>29</sub>H<sub>60</sub> | 0.99±0.12 |
| C30 | C<sub>30</sub>H<sub>62</sub> | 0.35±0.12 |

## Notes

### Competing Interest Statement

The authors have declared no competing interest.

